# High-Resolution Molecular-Dynamics Simulations of the Pyruvate Kinase Muscle Isoform 1 and 2 (PKM1/2)

**DOI:** 10.1101/2024.01.07.574528

**Authors:** Quentin Delobelle, Théo Jaffrelot Inizan, Olivier Adjoua, Louis Lagardère, Frédéric Célerse, Vincent Maréchal, Jean-Philip Piquemal

## Abstract

Glucose metabolism plays a pivotal role in physiological processes and cancer growth. The final stage of glycolysis, converting phosphoenolpyruvate (PEP) into pyruvate, is catalyzed by the pyruvate kinase (PK) enzyme. Whereas PKM1 is mainly expressed in cells with high energy requirements, PKM2 is preferentially expressed in proliferating cells, including tumor cells. Structural analysis of PKM1 and PKM2 is essential to design new molecules with antitumoral activity. To understand their structural dynamics, we performed extensive high-resolution molecular dynamics (MD) simulations using adaptive sampling techniques coupled to the polarizable AMOEBA force field. Performing more than 6 µs of simulation, we considered all oligomerization states of PKM2 and propose structural insights for PKM1 to further study the PKM2-specific allostery. We focused on key sites including the active site and the natural substrate Fructose Bi-Phosphate (FBP) fixation pocket. Additionally, we present the first MD simulation of biologically active PKM1 and uncover important similarities with its PKM2 counterpart bound to FBP. We also analysed TEPP-46’s fixation, a pharmacological activator binding a different pocket, on PKM2 and highlighted the structural differences and similarities compared to PKM2 bound to FBP. Finally, we determined potential new cryptic pockets specific to PKM2 for drug targeting.

**Entry for the Table of Contents:** 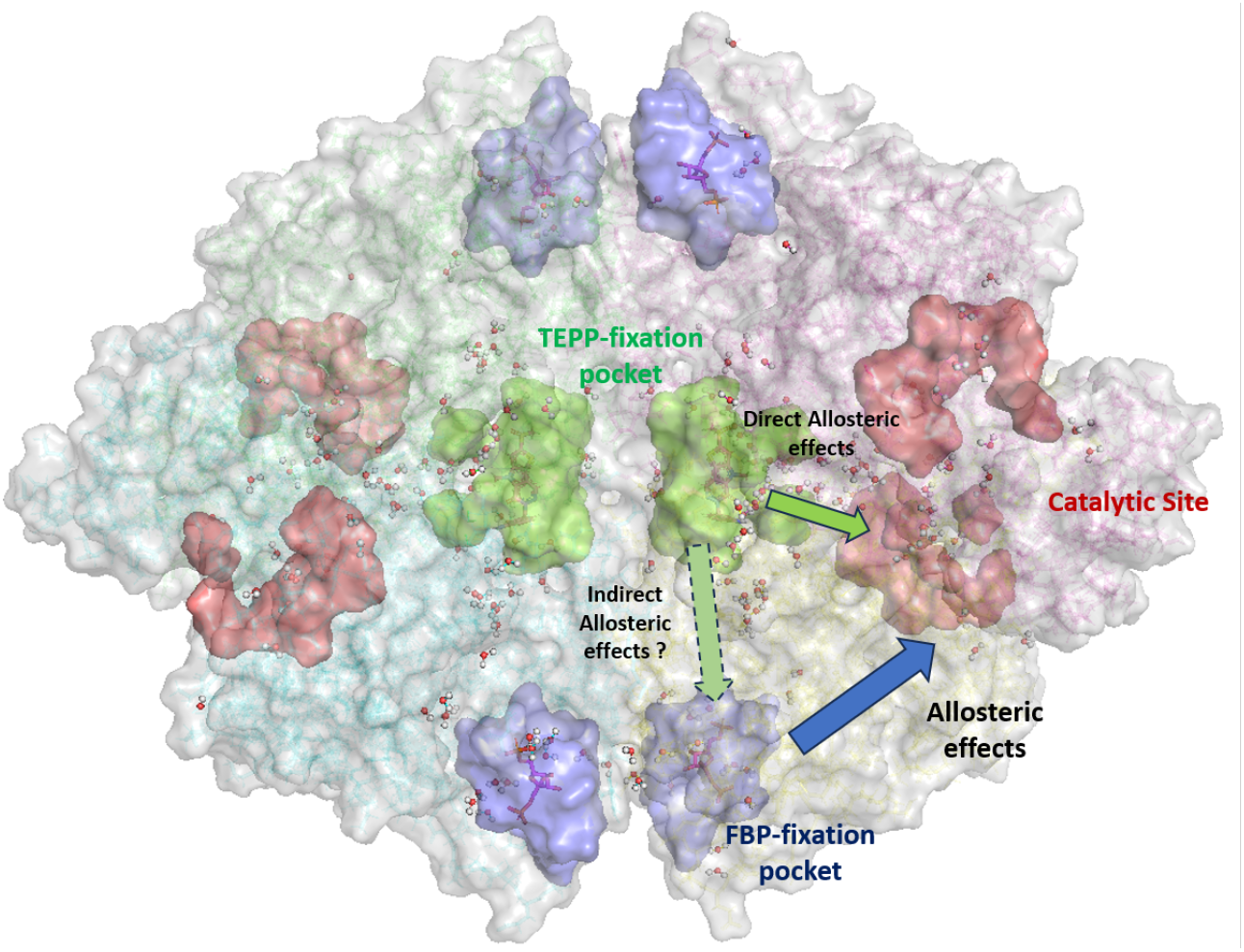

Producing more than 6*µ*s of cumulated simulation time with the AMOEBA polarizable force field, we determined key structural properties of the PKM2 (catalytic, FBP and TEPP) and PKM1 enzymes binding sites to determine new cryptic pockets for further antiviral/antitumoral drug design.

## Introduction

In normal mammalian cells, ATP is produced by both glycolysis and mitochondrial oxidative phosphorylation (OX-PHOS). However, in most tumor cells ATP is produced mainly by glycolysis, even in the presence of oxygen ^[1]^. One of the advantages of preferential activation of glycolysis is the accumulation of metabolic intermediates useful for tumor cell proliferation, such as molecules required for the biosynthesis of nucleic acids, proteins, lipids, and lactate required to balance the NAD+/NADH redox balance ^[2]^.

Glycolysis is a metabolic process encompassing 10 different steps^[3]^. It produces two molecules of ATP (adenosine triphosphate), two molecules of pyruvate and two molecules of NADH (Nicotinamide Adenine Dinucleotide) per molecule of glucose. The final stage of glycolysis is a rate-limiting and irreversible phase that converts phosphoenolpyruvate (PEP) to pyruvate. This reaction is catalyzed by pyruvate kinase (PK). There are 4 tissue-specific isoforms in mammals: PKL (Pyruvate Kinase Liver-type) is expressed in the liver, kidney, and intestine tissues, and PKR (Pyruvate Kinase in Red-blood cells) in erythrocytes ^[4]^. Whereas PKM1 (Pyruvate Kinase Muscle isoform 1) is expressed in tissues with high energy requirements such as brain, heart, and muscle tissues ^[5]^, PKM2 (Pyruvate Kinase Muscle isoform 2) is preferentially expressed in proliferating cells such as embryonic cells and tumor cells ^[6]^. These two isoforms are encoded from the same gene, PKM, which contains 12 exons by alternative splicing generating PKM1 (exon 9) or PKM2 (exon 10) ^[7,8]^. PKM1 and PKM2 encompass a total of 531 amino acids while differing by 22 of them ^[9]^.

PKM1 is a constitutive tetrameric active enzyme with a high affinity for PEP. Conversely, PKM2 exists in three different forms depending on the cellular compartments: an inactive monomer, a nuclear dimer associated with a low enzymatic activity and a cytoplasmic active tetrameric form. Following the binding of fructose-1,6-biphosphate (FBP) on cytoplasmic PKM2 (either in the dimeric or tetrameric state), the allosteric activation of PKM2 promotes its tetrameric form.^[10]^. Additionally, the dissociation of the tetramer favors the accumulation of the PKM2 dimer, which exhibits pro-tumor activity ^[11]^. The PKM2 tetramer itself is also known to oscillate between two structural organizations: a less active tight (T) form and a fully activated relaxed (R) form. The relative prevalence of these forms is directly associated to the PK immediate environment and the presence of regulating metabolites. PKM1 and PKM2 structural differences may explain why PKM1 is not regulated by allosteric activator ^[12]^. On the opposite, PKM2 enzymatic activity is highly regulated by many endogenous metabolites. It is important to point out that such metabolites can either have the role of an activator (for example: serine ^[13]^, SAICAR (Succinylaminoimidazole-carboxamide riboside ^[14]^) and FBP ^[15]^) or the role of an inhibitor (for example tryptophan, the T3 thyroid hormone ^[16]^ or HMGB1 ^[17]^ for example but as another protein). Each monomer of PKM2 contains 3 domains (A/B/C), a short N-terminal region, and an active site. The A domain constitutes the main part of the protein (from residues 44-116 to 219-389). This region is important since the dimeric form of PKM2 is formed through the weak bonding of two of its monomers interacting with this region. The B domain is characterized by an external region formed by residues 117-218. It constitutes a part of the active site of PKM2 (PEP fixation) which is structured by an interface between A and B domains. Finally, 44 of the 56 amino acids encoded by PKM2-specific exon 10 are located in the C domain, which encompasses the allosteric binding site and stabilizes the quaternary structure of the tetrameric form of PKM2. The N-terminal region is localized on residues 1-43 of PKM2 ^[18]^. PKM2’s central role in tumor processes makes it a prime target for the design of new anti-tumor molecules and several drugs have been proposed to modulate PKM2 activity and oligomeric state. They act as an activator promoting the active R-form (TEPP-46, DASA-58 ^[19]^). Consequently, it is essential to have a better understanding of the structural dynamics of PKM1 and PKM2 and to identify the conserved sites that could lead to the design of new antitumor molecules. Notably, it is especially relevant to investigate the structure of the PKM1 isoform to understand its main structural differences compared to PKM2 to propose molecules capable of promoting the conversion of PKM2 from the pro-tumor dimer into an active tetramer.

In this study, we provide all-atom high-resolution molecular dynamics simulations of both PKM1 and PKM2 enzymes and propose a comparative analysis of their key residues and functional sites. In particular, our study focuses on the characterization of the FBP-binding pocket and on the PEP active site which are both closely related to the specificity of activity of both proteins. To provide high-resolution simulations approaching realistic biological timescales, we use adaptive sampling molecular dynamics coupled to the AMOEBA polarizable force field ^[20,21]^. Indeed, electronic polarization plays a critical role in defining a protein’s conformational space by altering the stability of secondary and quaternary protein structures and solvation ^[22–25]^. Thanks to such methodology coupled to the use of the GPU-accelerated Tinker-HP molecular dynamics package ^[26,27]^ (GPU=-Graphics Processing Unit), it is possible to simulate the weak interaction profiles ^[28–31]^ of these proteins. We therefore studied PKM1 and PKM2 in their different oligomeric states, notably in the presence or absence of their known effectors.

## Results and Discussion

In this section, we analyzed the structural dynamics of PKM1 and PKM2. To do so, the alpha carbon backbone of each protein residues was chosen to make these measurements in order to obtain an interpretation of the arrangement of the PKM proteins.

### Understanding PKM1/2 organization during the oligomerization process

First, we compared the structural variations between PKM1 and the different oligomerization states of PKM2 by analyzing the Root-Mean Square Deviation (RMSD) and the Radius of Gyration (RG). The results revealed an increasing correlation of RMSD and RG as PKM2 oligomerization complexified. Starting with the PKM2 monomer (see Figure 1A) we observed a specific region of large variations of RMSD and RG (2-7 Å and 24.75-26.50 Å respectively) during the simulation. High densities of structures were found at RMSD 3.6 Å and RG 25.5 Å. Overall, the monomeric state was characterized by an important variability as suggested by the absence of correlation between RMSD and RG. Since the simulations enabled us to explore vast conformational spaces, this indicated that the monomer was poorly structured. In contrast, the dimer displayed a narrower distribution of RMSD around 2.5-5.5 Å (see Figure 1B) was observed with some larger variations of RG at 31-40 Å. Overall, a single high density of structures is observed with a significant correlation between RMSD and RG, a sign of the superior structuring of the PKM2 dimer compared to its monomer. In the case of the apo PKM2 tetramer (see Figure 1C), we observed a larger RMSD range compared to the dimer (2.2-7.4 Å) which correlated with higher RG values (40.7-45.5 Å). Again, a concentration of high densities was found but an almost linear RMSD/RG correlation was observed. This observation was even more striking when the PKM2 tetramer was bound to FBP (see Figure 1D). Indeed, FBP appeared to have a stabilizing effect on the enzyme structural variability as indicated by the more restricted distribution of RMSD values (from 1.7 to 5.2 Å). Again, an extremely dense region of structures (3.5 density at 3.75 Å RMSD and 43.75 Å RG) was found, as sign of a fully structured enzyme.

**Figure 1.**
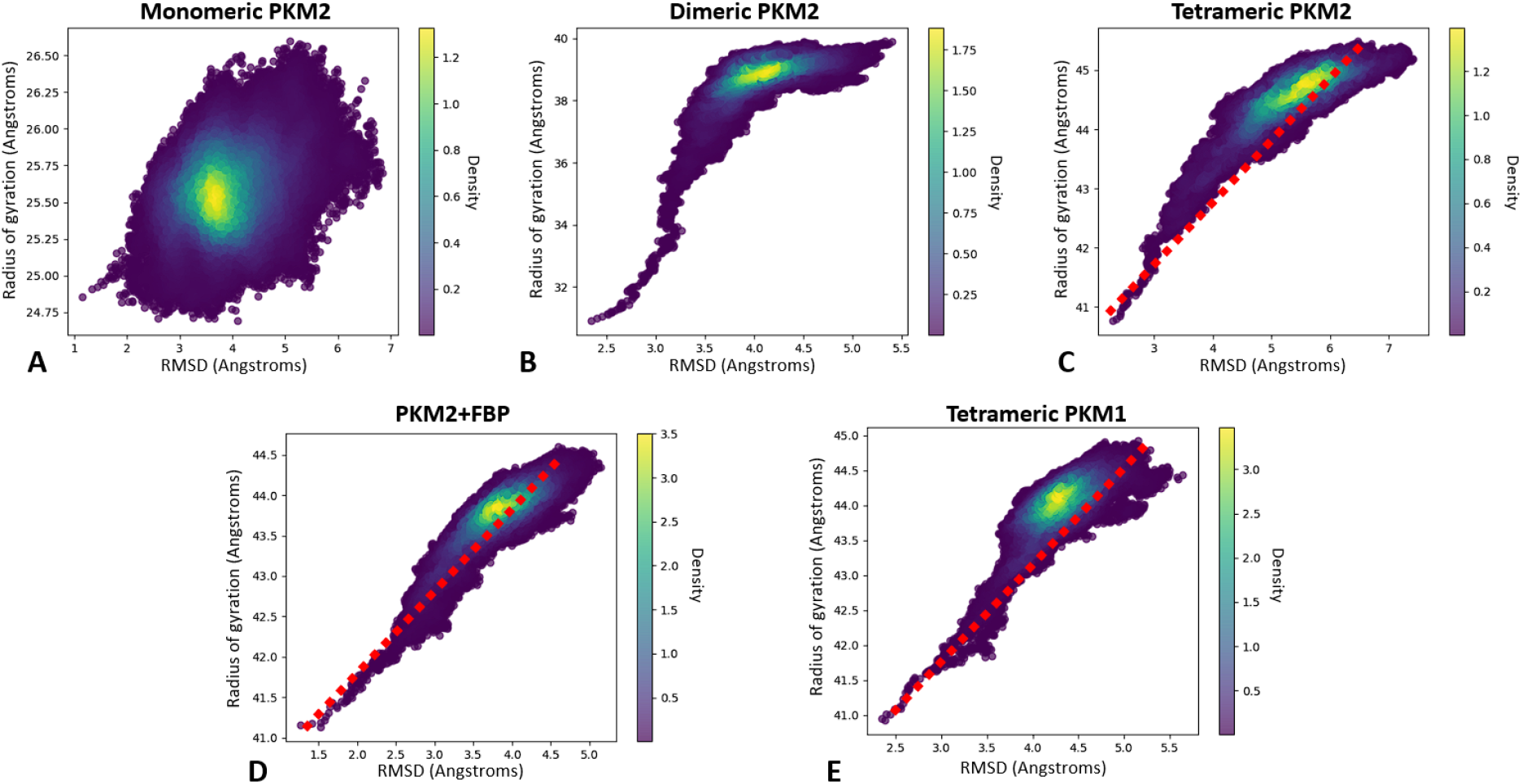
The global analysis of the RMSD vs RG shows the structural changes of PKM2 during the oligomerization process. We also analyze PKM1/2 profile similarities. (A-D) Graphical representation of the RMSD/RG analysis for each states of PKM2 molecular dynamics simulation : monomer **(A)**, dimer **(B)** and free-tetramer **(C)** or associated to FBP **(D), (E)** Same representation for the tetrameric form of PKM1.

We then compared the different PKM2 structures to the PKM1 tetramer. As shown in Figure 1E, the PKM1 profile appeared to be close to the tetrameric PKM2 when associated to FBP. Indeed, an almost identical dense region of structures has been observed in PKM1 compared to PKM2-FBP (3.4 density at 4.25 Å RMSD and 44 Å RG) as the RMSD/RG correlation was almost linear.

### PKM1/2: Analysis of residues and domains stability during the MD simulations

The next step investigated key amino acids present in both PKM isoforms as a structural markers of the flexibility of specific domains. To do so, we studied the Root-Mean Square Fluctuation (RMSF) to quantify the mean deviation for each residue throughout the simulations by comparison them to their position in the PDB reference. The first immediate conclusion linked the enzymes’ stability to the oligomerization process. Indeed, in the case of the PKM2 monomer (see Figure 2A), our results showed an important flexibility affecting the entirety of the structure. The greatest flexibility was observed in the B-domain (residues 116-217), where RMSF values reached higher than 4 Å, indicating significant variability. We also observed a large flexibility for three large regions on the A1-domain (residues 45-115) and on additional residues (residues 290-425 and 475-525) that composed a large part of the A2-and C-domains. A structural stabilization was observed for the dimeric state of PKM2 (see Figure 2B) as the A2-domain (residues 290-400) appears to have lost most of its flexibility compared to the monomer. However, the dimer still retains some variability in its structural dynamics located at the A1 (residues 50-107) and C-domains (residues 400-457 and 470-531),contrary to the apo PKM2 tetramer known for better enzyme activities (see Figure 2C). Indeed, in this state the A1-and C-domains residues lost most of their structural variability. A few conserved pockets exhibiting some flexibilities were found around residues 30-110 and 470-531. Such residues participate to the structure of the active site and to the FBP-fixation pocket of PKM2.

**Figure 2.**
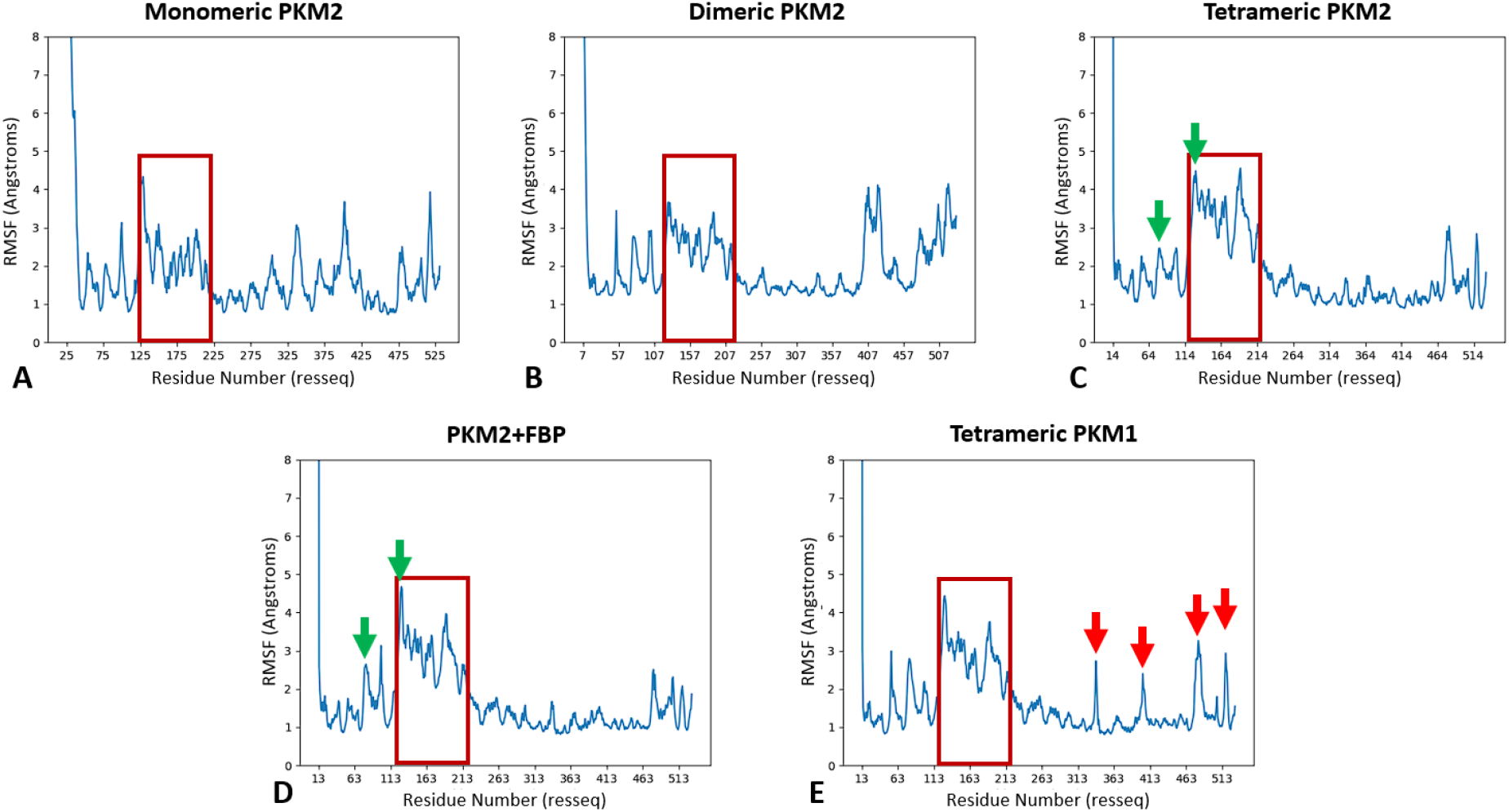
The RMSF analysis shows PKM2’s structuration during the oligomerization and FBP-fixation processes and confirms the tetrameric PKM1/2 analogies on key domains. (A-D) Graphical representation of the RMSF analysis for each state of PKM2 molecular dynamics: monomer **(A)**, dimer **(B)** and free-tetramer **(C)** or associated to FBP **(D), (E)** Same representation for the tetrameric form of PKM1. Here the PKM2 domains contain the following residues : 44-115 for A1, **116-217 for B (red square)**, 218-388 for A2 and 389-531 for C-domain. Local differences between apo and holo PKM2 forms bound to FBP (green arrows), and between holo form of PKM2 bound to FBP and PKM1 respectively (red arrows) are indicated.

Considering its central role in the making of the active site, it is also important to note that each oligomeric state conserved important variability in the B-domain. We then evaluated the allosteric effect of the fixation of FBP on the tetrameric PKM2 (see Figure 2D). If our results showed global RMSF profiles that are extremely close between the holo and apo systems, local differences were observed. First, the FBP fixation pocket was less flexible in the holo-FBP state compared to the apo state localized at residues 470-531. Second, a relative flexibility for A1 (residues 70-110) and certain residues of the B-domain located at 75-80 and 115-120 was still observed, these residues corresponding to key amino acids involved in the active site (see green arrows, Figure 2C).

Our results showed a close structural homology between the PKM1 tetramer (see Figure 2E) and the PKM2 holo form bound to FBP, notably in the B-domain. However, some specific regions remain different. Some couples of residues (residues 325-335, 390-405, 470-490, 515-525) present a relatively high flexibility (see red arrows, Figure 2E). Altogether, these results gave a full picture of the local structural effects of the progressive structuring of PKM2 during the oligomerization process. It confirmed the strong structural homologies between PKM2-FBP and PKM1.

### Structuration of the active site and oligomerization states of PKM2

We next focused on the PKM2 key reactivity sites and started by analyzing the enzyme active site. To do so, the distances between key couples of residues defining structural markers were studied to evaluate the flexibility of the B-domain. We measured the distance between residue 53 of the A-domain and residue 129 from the B-domain, both located at the external part of the active site. They were chosen to characterize the global motion of the B-domain. To study the active site structural organization, we monitored the distance between residues 178 and 296 (see Figure 3E) localized at the junction between the A- and B-domains. While the monomeric state (see Figure 3A and 3C, red curve) presented a restricted distance between residues 53 and 129 (around 20.5 Å), a bimodal distribution was observed between residues 178 and 296. Such distribution, presented in Figure 4 for the monomeric state of PKM2 was present in the other PKM2 oligomers. Looking at the dimer (see Figure 3A and 3C, grey curve), we observed a profile similar to the monomer for the distance between residues 53 and 129. A different bimodal distribution was observed for the distance between residues 178 and 296 with values centered at 5 and 7.5 Å, which indicates a more constrained spatial extension of the active site (**see Figure S1 in Supplementary Informations**). Then we finally considered the PKM2 tetramer in its apo state (i.e. without its natural ligand, see Figure 3A and 3C, green curve). Here, we noticed a higher distance between residues 53 and 129 coupled to a distance between residues 178 and 296 of 8-8.5 Å. Such a result indicated a higher flexibility of the B-domain that can be linked to a stronger structuration of the PKM2 active site. This observation aligns with the documented activities of the various oligomers ^[10]^. When PKM2 was associated with FBP, i.e. forming the holo state (see Figure 3B and 3D, yellow curve), we observed a higher distance between residues 53 and 129 compared to the apo state. The distance between residues 178 and 296 was similar to the apo tetramer but with a slightly narrower distribution centered at 8.25 Å. This was in line with the known stability of the PKM2 tetramer when bound to FBP ^[15]^.

**Figure 3.**
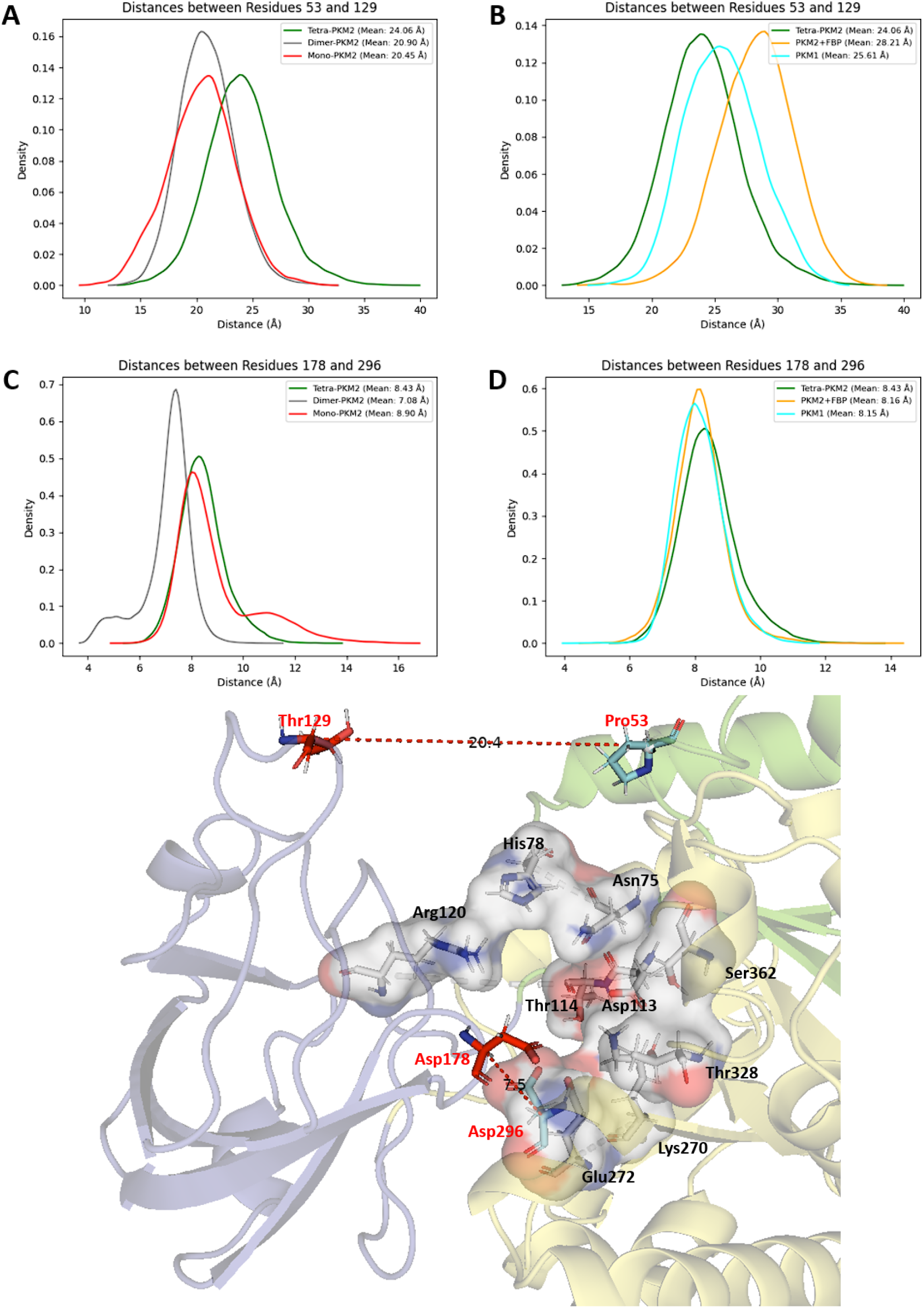
The configuration of the active site is linked to the oligomeric conformation of PKM1/2. (A-B) Graphical representation of the distance between residues 53 and 128 (in the active site) between each state of PKM2 within the simulations with the monomer, dimer and tetramer-apo **(A)** or between tetramer-apo PKM2, holo PKM2 with FBP and also the tetrameric state of PKM1 **(B), (C-D)** Same representation for residues 178 and 296 between each state of PKM2 **(C)** or PKM2 and PKM1 **(D). (E)** Representation of all the catalytic residues composing the active site with the selected residues highlighted in red.

**Figure 4.**
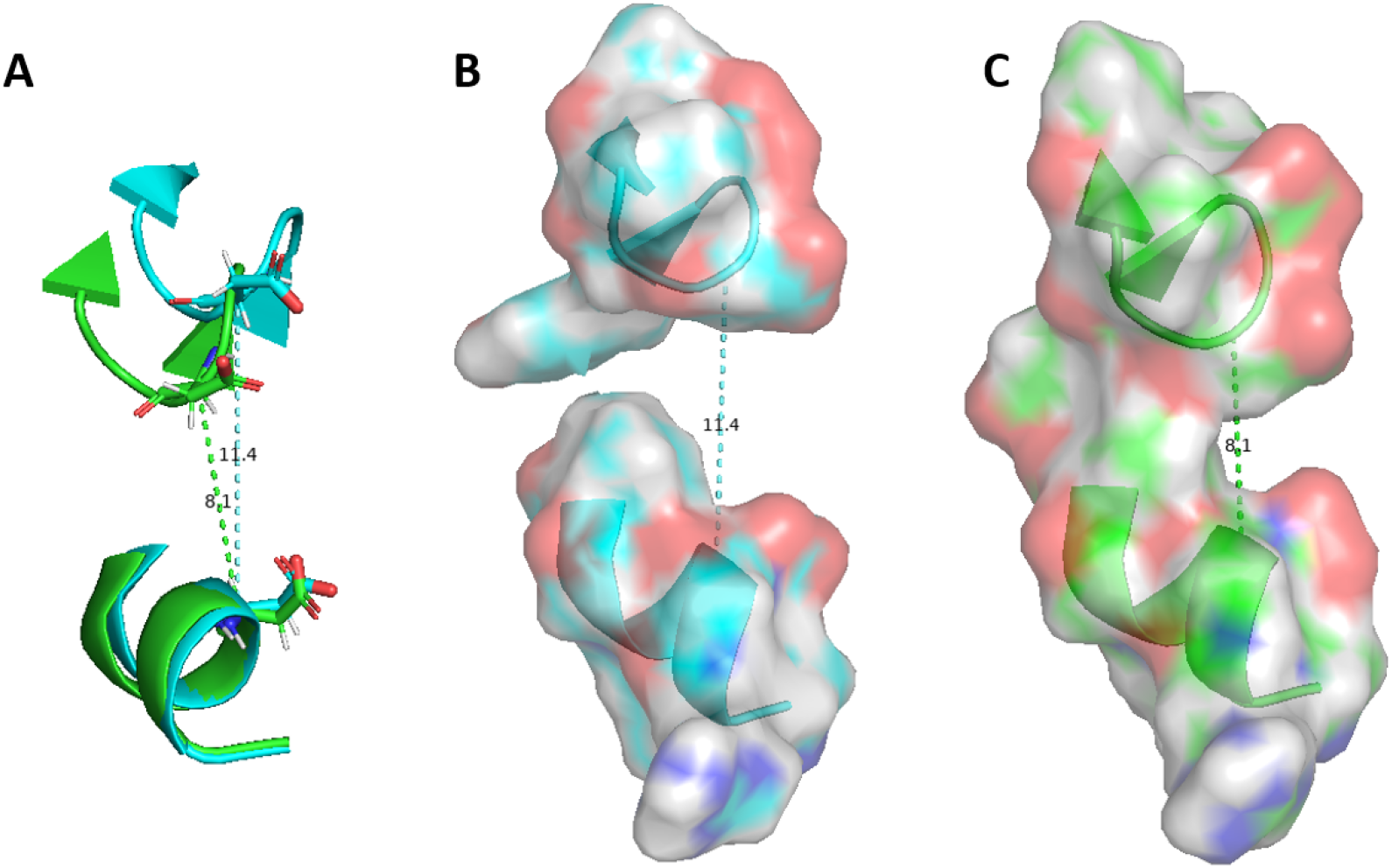
Comparative study of different states of the active side internal part inside monomeric PKM2. (A) Graphical representation of the distance between residues 178 and 296 for two different frames of the bimodal distribution, **(B)** Surface model showing the high-distance state (*≈* 11.5 Å) **(C)** or the more enclosed one (*≈* 8 Å). Three amino acids before and after residues 178 and 296 are represented to obtain a better surface model.

Finally, we characterized the active site in the tetrameric state of the PKM1 protein to establish a comparison with its PKM2 counterpart (see Figure 3B and 3D, cyan curve). The observed distance profiles were found to be intermediate between the values observed from the apo and holo-FBP state of the PKM2 tetramer. Concerning the external part of the active site, the distance between residues 53 and 129 was lower compared to holo-PKM2 while remaining higher than the distance measured for apo-PKM2, which indicated an higher-level of structural organization for the PKM1 active site compared to the free-PKM2. Along the same line, the distance between residues 178 and 296 exhibited a very similar profile compared to the holo-PKM2 structure (i.e. mean of 8.15 Å vs 8.16 Å for the free-tetrameric-PKM2 state). Overall, our data showed a net effect of the FBP allosteric effector on tetramer active site structuring that appeared relatively similar to the PKM1 isoform.

### Structural Dynamics of the FBP-fixation pocket : Allosteric and Oligomeric modulations

Here, we compared the structural organization of the region containing the FBP-fixation site of PKM2 with the homologous region in PKM1. The distance between residue 433 and 518 has been studied to evaluate the global flexibility of the FBP-fixation site since these residues are located at the extrema of the pocket (**see Figure S2 in Supplementary Informations**). We first compared each possible PKM2 oligomeric states used for our simulations. The results obtained for the monomer (see Figure 5A, red curve) showed a highly variable profile of distances with a large density of structures centered at 7.5 Å that were also coupled with a very large proportion of higher distances. This variability suggests that the FBP-fixation pocket is not well-formed or neither stable in the monomeric state of PKM2. Considering the PKM2 dimeric form (see Figure 5A, grey curve), we observed higher distances compared to the monomeric state. More precisely, a unique and wide distribution was found with distances ranging from 13 to 21 Å which was not in favor of an overall well-structured FBP-fixation pocket. A bimodal distribution was observed in the apo PKM2 (see Figure 5A, green curve), with a more preponderant population of structures centered at 12.5 Å and a second one at 9.5 Å. This particular distribution is a signature for the co-existence of two distinct states for the tetrameric PKM2. For PKM2 bound to FBP, we noticed a complete inversion of the observed 2-states distribution compared to the apo-PKM2 state (see Figure 5B, yellow curve). In this case, a major population of structures was centered around a mean distance at *≈* 7 Å whereas a second population was center at *≈* 8.5 Å. These results confirmed that FBP played a major role in the structuring of the FBP fixation pocket.

**Figure 5.**
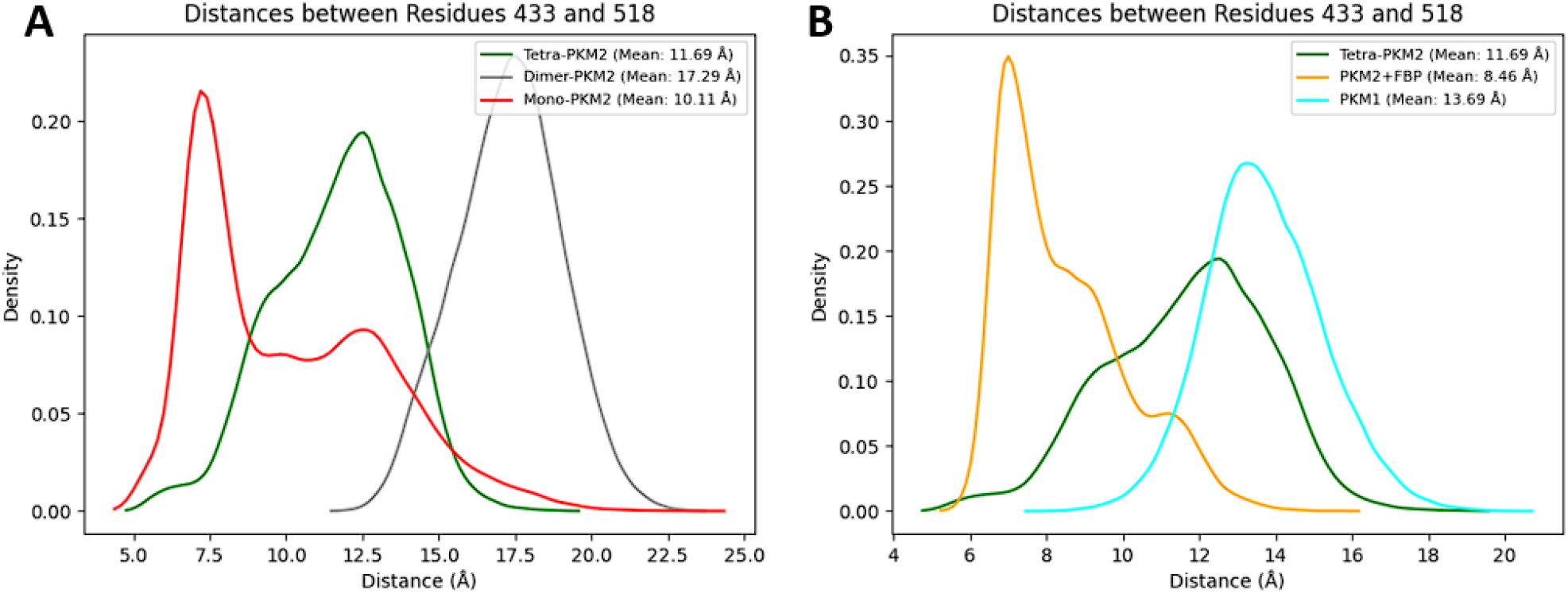
Global structuring of the FBP-fixation pocket during PKM2 oligomerization and FBP-docking process and comparison with PKM1 possible fixation site. (A) Graphical representation of the distance between residues 433 and 518 (localized at the FBP-allosteric pocket) for each oligomeric state of PKM2 molecular dynamics, **(B)** Same representation for the apo and holo PKM2 tetrameric forms and the tetrameric form of PKM1.

We then compared the FBP-binding region of PKM2 to its PKM1 counterpart. The data obtained from the tetrameric state of PKM1 (see Figure 5B, cyan curve) showed a distribution of the distance centered at 13.7 Å, a much higher distance than the one observed in apo-PKM2 and in holo-FBP PKM2. This data confirmed that the putative allosteric site in PKM1 did not appear to be structured enough to have a viable fixation pocket able to bound FBP. Examples of the structural comparison between the free and FBP-bound PKM2 and PKM1 can be found in Figure 6 and **Figures S3 and S4 of the Supplementary Informations**. These models showed on one hand the open pocket in the case of the tetrameric state of PKM1 (with a loop orientated in the external side) but also the importance of FBP to structure such PKM2 region since its absence let loose the regulatory loop necessary to the pocket stability.

**Figure 6.**
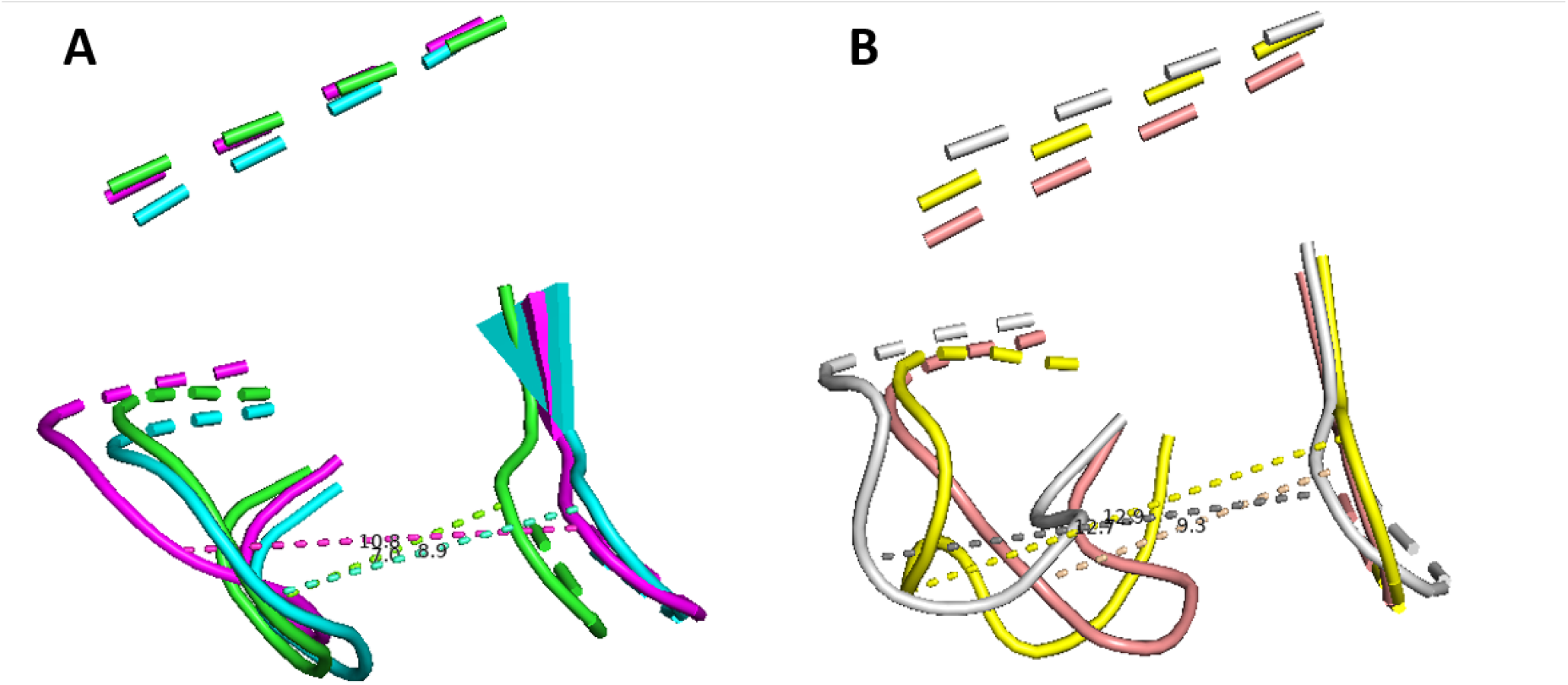
Comparative study of different states of the FBP-fixation pocket for PKM1 and PKM2. (A) Cartoon model of three frames aligned during simulation of PKM2 bound to FBP highlighting the three major states at *≈* 7*/*9*/*11 Å in comparison to **(B)** two frames obtained from the simulation of free-tetramer of PKM2 (in light pink and grey, highlighting the two states at *≈* 9*/*13 Å) and a single state of PKM1 in yellow at *≈* 13 Å.

### PKM2 associated with the TEPP-46 pharmacological activator

For our next analysis, we studied the structural dynamics of PKM2 associated with TEPP-46, a pharmacological activator of the PKM2 tetramerization. TEPP-46 binds at the interface of two monomers through their A-domains (**see Figure S5 in Supplementary Informations**, red zone), an interaction pocket different from the one present with FBP **(see Figure S5 in Supplementary Informations, orange frame)**. Such differences also impact the stoichiometry of TEPP-46 compared to FBP since 4 molecules of FBP can bind the PKM2 tetramer, while only two molecules of TEPP-46 can bind it at the same time. First, the RMSF data (see Figure 7A) gave a similar profile to the one observed for PKM2-FBP with minor local differences (residues 390-405 and 470-490), that were located in the areas of interaction with TEPP-46 and in some residues for FBP-fixation pocket respectively. As for the studies of PKM2 tetramers, the B-domain (residues 116-217) retained the same flexibility profile as the one observed in the presence of FBP. To gain additional structural information, we studied in more detail the PKM2 active site (see Figure 7B). For the distance between residues 53 and 129, similarities were noted between PKM2 bound to FBP and PKM2 linked to TEPP-46. The data showed that the distance at the A-B junction (residues 178 and 296) was very similar to the one observed when PKM2 was linked to FBP. Conversely, the fixation of TEPP-46 had an opposite effect on the movement of the B-domain. Indeed, the distance between amino acids 53 and 129 was reduced and exhibited a bimodal profile at 12 and 19 Å indicative of a closed proximity between the B- and the A-domain. Altogether, this was consistent with an improved enzymatic activity for TEPP-46 bound to PKM2 compared to PKM2 tetrameric-free state. Finally, the structural analysis of the FBP-pocket for the PKM2 tetrameric state bound to TEPP-46 (see Figure 7C) showed that the distance between residues 433 and 518 was extremely close (i.e. centered at *≈* 7.5 Å) compared to the one present in the FBP-PKM2 structure. However, in the case of PKM2 bounded to TEPP-46, no ligand was localized in this space since the TEPP-46 fixation pocket is located in a different region of PKM2.

All the important sites for PKM2 are presented in **Figure S8 in Supplementary Informations**, including additional structural views of the TEPP-46 fixation site.

**Figure 7.**
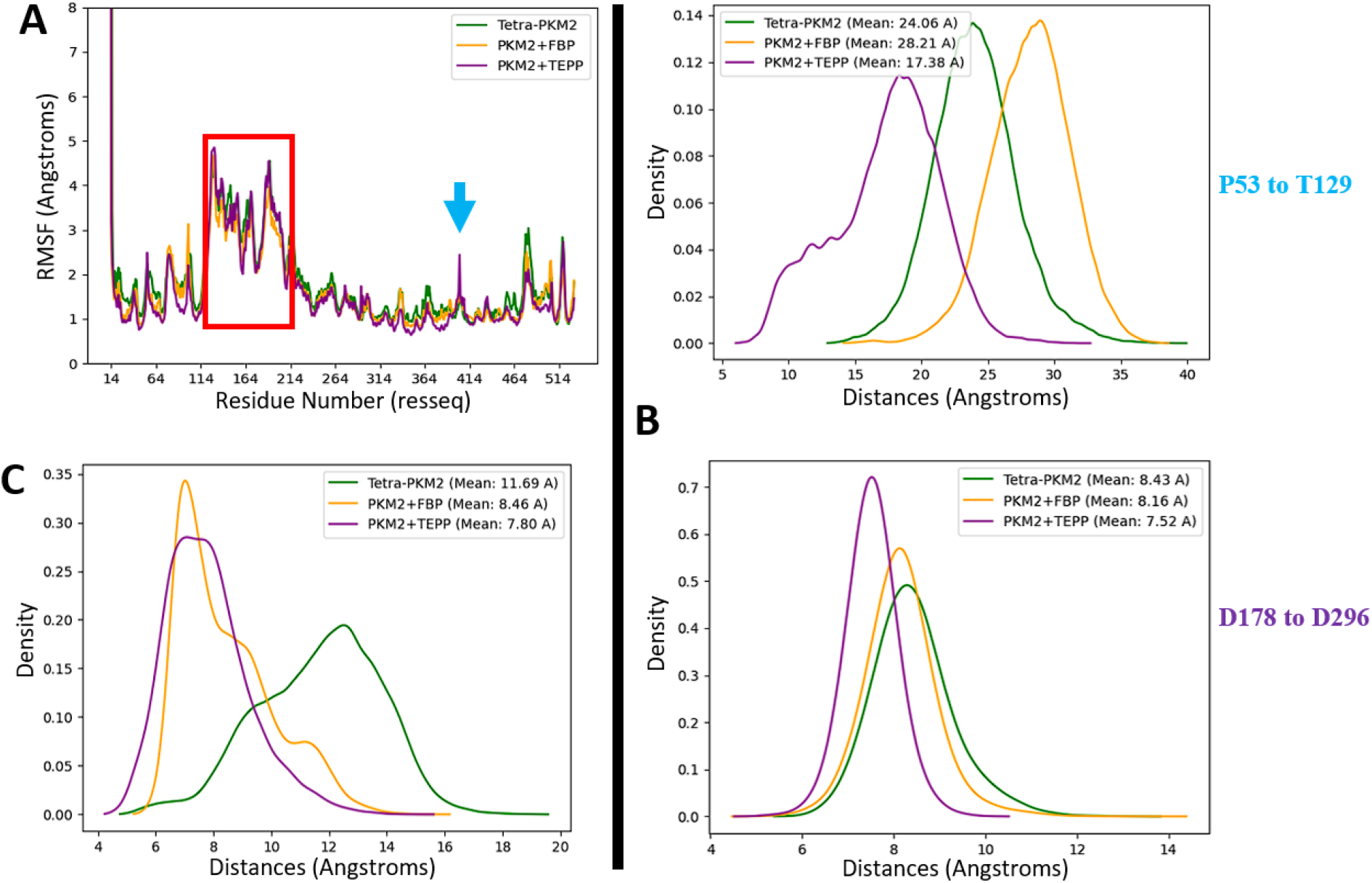
Structural data associated to the TEPP-46 fixation on a tetrameric PKM2. (A) Graphical representation of the RMSF analysis, **(B)** kernel density of the distance between residues 53 and 128 (in the active site) and **(C)** distance between residues 433 and 518 (localized at the FBP-allosteric pocket) for different tetrameric state of PKM2 with or without natural or pharmacological effectors.

### Comparative analysis of druggability: searching for PKM2-specific cryptic pockets

We used the presented structural models to identify new cryptic pockets. Such pockets are three-dimensional structures that are generally hidden or partially accessible and that could be targeted by new pharmacological modulators. Given its importance in tumor pathologies, we wanted to identify pockets specific to the tetrameric form of PKM2. We first searched for these pockets using the crystallographic structures of the PKM1 tetramer (3SRF) and of the FBP-bound PKM2 tetramer (3SRD). The search was conducted using the DoGSite Scorer software, an automated pocket detection and descriptor calculation tool ^[32]^. 63 and 50 pockets were detected for PKM1 and PKM2 respectively. Several known pockets were found including the active site, the binding pocket for regulatory amino acids (Ser, Phe), the FBP binding pocket and the TEPP-46 binding site (Figure 8). To determine new pockets that were specific to PKM2 and in order to avoid potential adverse effects targeting both forms, we have eliminated the pockets which were common to both PKM1 and PKM2. After such an elimination, 15 and 13 pockets were retained for PKM1 (called M1-X, with X representing the pocket number in the DoGSite Scorer software) and PKM2 (M2-X) respectively. These pockets, identified in the crystallographic structures, were used as a reference for the remainder of this analysis. For each tetramer, PKM1 and PKM2-FBP, 8 structures derived from molecular simulations (Tinker-HP) were randomly selected and analyzed by DoGSite Scorer. The identified pockets were then compared with the reference pockets. The comparison was made by analyzing the residues of all pockets and assigning them to the reference pocket with the most common number of overlapping residues. This pocket overlap criterion is based on a maximum number of common residues, set at 5, but also on the addition of an overlap ratio value corresponding to the ratio between this maximum number of residues and the total number of residues corresponding to the predicted pocket, set at 0.25 (i.e. less than 25% total sequence identity). These pockets were named in the same way as the reference pockets T_M1-X or T_M2-X, (T corresponding to the structures derived from Tinker-HP molecular simulations). The results of this analysis are displayed in Figure 9. The known pockets - active site and amino acid binding sites, FBP and TEPP-46 - were detected in all sampled structures. It should be noted that the presumed FBP binding pocket on PKM1 was found in all the sampled structures, even though previous results suggest that it is not functional. The different pockets detected from the two initial crystal structures, are detected either (1) in a large number of the sampled structures (i.e. ‘M2-0’, ‘M2-2’, ‘M1-0’ or ‘M1-33’), or (2) in a few of them, such as ‘M2-37’ or ‘M1-55’. This diversity validates the use of molecular simulations with adaptive sampling, but it should be pointed out that none of these initial pockets containing residues specific to PKM1 and PKM2 exons is isoform-specific. This strategy identified 11 new cryptic relatives on PKM2. Two pockets are of particular interest (‘T_M2-8’ and ‘T_M2-11’) as they are specific to the PKM2 tetramer bound to FBP. The ‘T_M2-8’ pocket occupies an average volume of 174 Å^3^ associated with an average druggability score ranging from 0.37 to 0.45 (based on pocket volume, hydrophobicity and accessibility) according to analyses performed with DogSite **(see detailed results on druggability score on Figure S6, Supplementary Informations)**. Pocket ‘T_M2-11’ occupies a volume of 126 Å^3^ with a druggability score of 0.25. Two other pockets, ‘T_M2-3’ and ‘T_M2-10’ were also more frequently detected for PKM2-FBP at 2 and 7 detections respectively. But they appear to be less specific, each detected on a single PKM1 structure (at an overlap ratio of 0.38 and 0.41 respectively). Nevertheless, their average volume and druggability score are indicated by their interest, particularly for ‘T_M2-10’ **(see Figure S7 in Supplementary Informations)**. Three of these pockets (not ‘T_M2-10’) are located at a crossroads between two chains of the PKM2 tetrameric structure. Furthermore, pockets ‘T_M2-3’ and ‘T_M2-11’ contain certain residues that appear to be involved in close interactions between chains within the tetramer, which could suggest that they are involved in the organization of the PKM2 tetramer. Targeting them could be an efficient lead to further characterize the interactions between the different chains of the tetrameric form of PKM2 and how the equilibrium between R- and T-state discussed in introduction is maintained.

**Figure 8.**
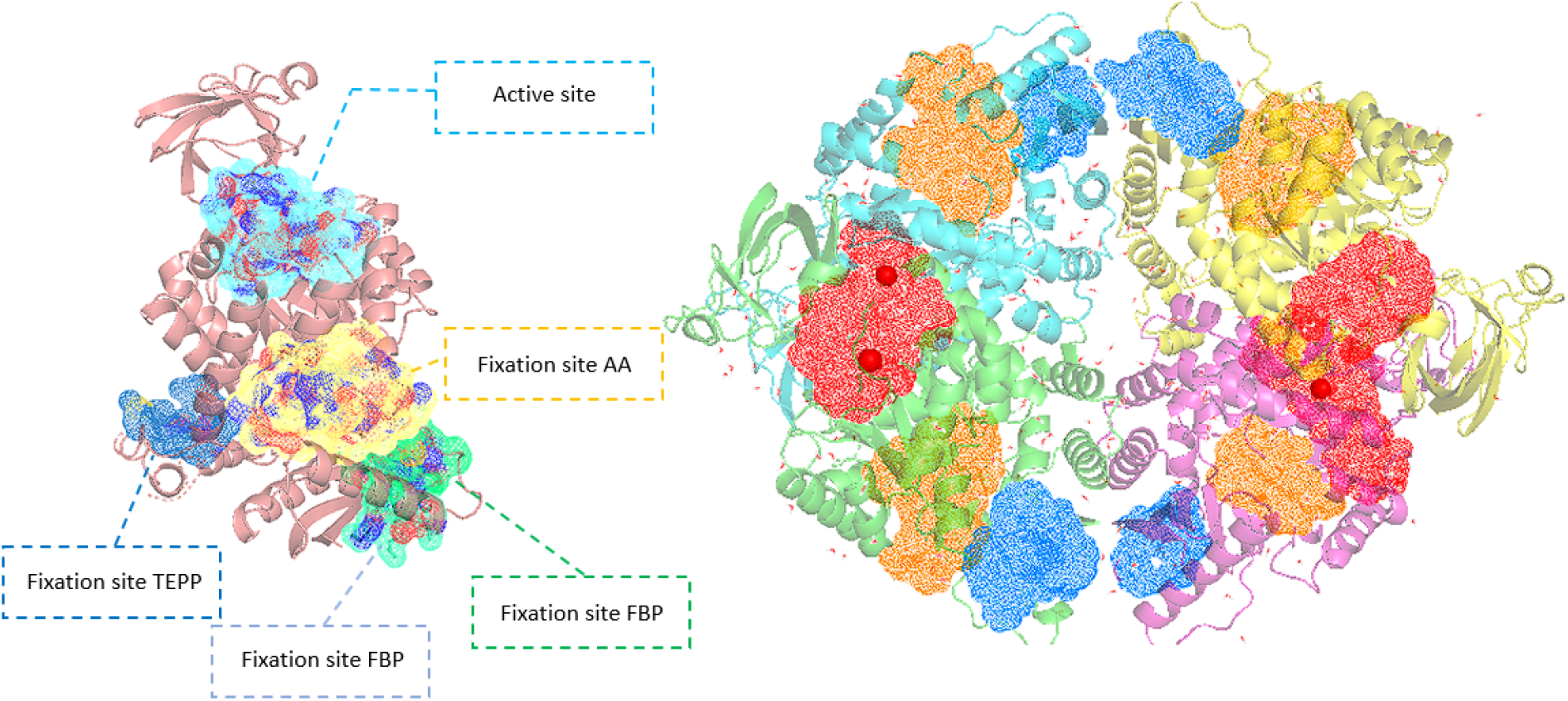
Representation of the major pocket locations for PKM1 (PDB:3SRF). One chain (left) and the tetrameric state (right) are both represented with their similar pockets except for the TEPP-46 pocket in the case of the tetramer since it is contained in a larger pocket. The active site, FBP and the amino acid fixation pocket are represented in red, blue and orange respectively in the case of the tetrameric representation.

**Figure 9.**
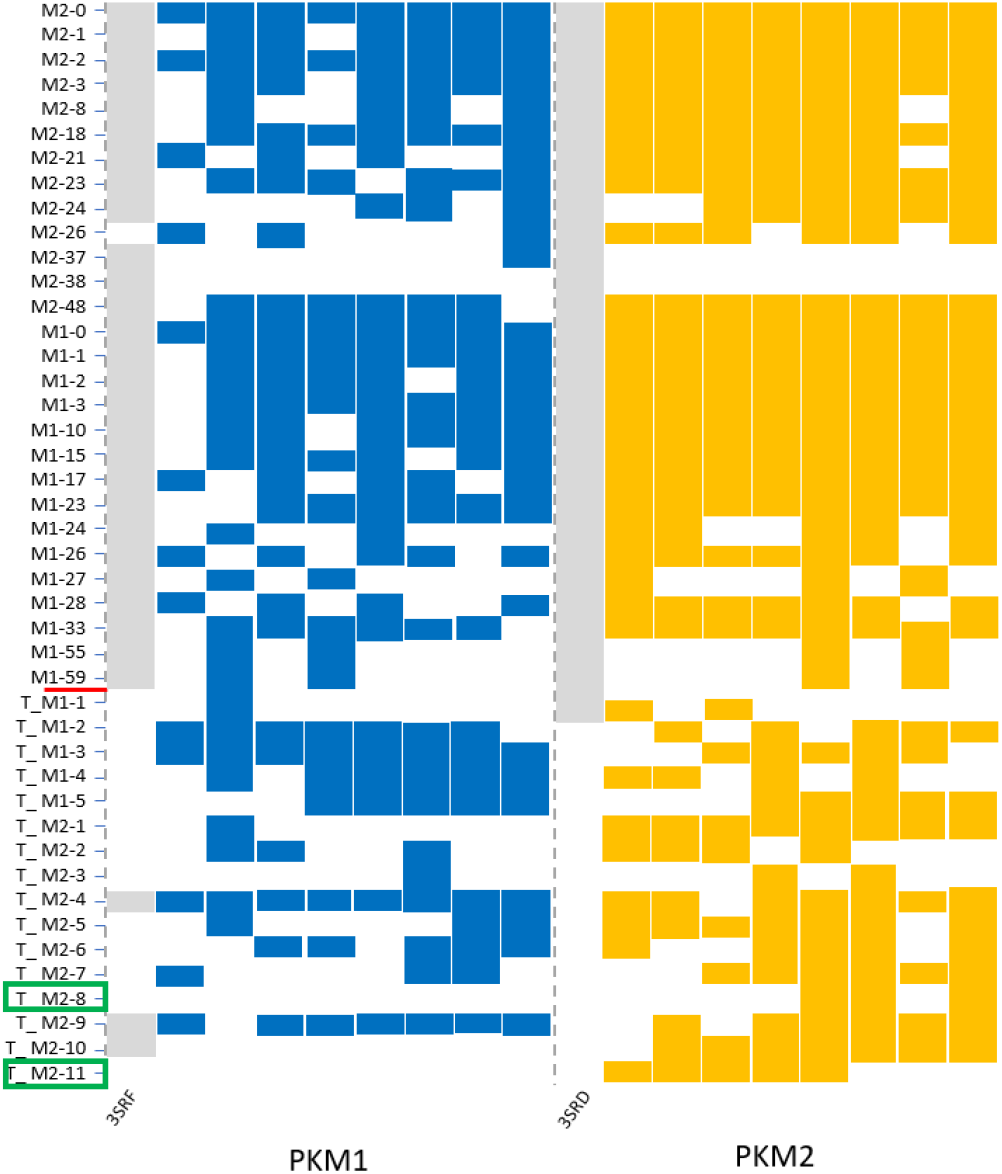
Schematic representation of pockets sampling detected via the DoGSite software. The pockets were detected from simulations based on the 3SRF (PKM1) and 3SRD crystal structures (PKM2) (in grey) associated to 8 additional structures extracted from Tinker-HP simulations for PKM1/PKM2, blue/orange respectively. The green rectangle highlights the pockets that were only detected for the PKM2 case.

## Discussion

We first compared the different oligomerization states of PKM2, ranging from monomer to tetramer and revealed an increasing stabilization of the protein in relation with higher oligomerization. We observed an increase in the linear correlation between RMSD and RG values, culminating in the tetrameric holo form. Such a correlation is therefore a good marker of the PKM2 conformational transition. These results are partially validated by the experimental data of Ponnusamy ^[33]^ that provided RG and RMSD results for a PDB structure of PKM2 that is similar to our simulations monomer data. Overall, our results clearly correlate with the experimentally observed differences in enzymatic activity ^[34]^. The dimeric form is less active whereas the tetrameric form of PKM2 exhibits a higher affinity for its PEP substrate and a better enzymatic activity. This enhanced activity across states strongly correlates with the progressive structuring of the active site. Additional information can be found directly by studying the evolution of the RMSF profiles: starting from an extremely vari-able monomeric state, the dimer stabilized itself while the tetrameric form of PKM2 bound to FBP exhibits the highest level of structural organization. Nevertheless, the B-domain (residues 116-217) still shows a high flexibility for each oligomeric state. This flexibility is reduced in the tetramer compared to the dimer, potentially explaining the higher substrate affinity observed in the tetramer ^[35]^. Further analysis of distances at key residues of the active site (53-129 and 178-296) supports this hypothesis. Indeed, the bimodal and more narrow value at the inside part of the active site of the dimeric form is extremely different from its tetramer counterpart. In the same line, the dynamically unstable internal junction at the active site of the monomer, which confirms the experimental data from Guo et al ^[36]^ who showed reduced enzymatic activity of the monomeric state. To conclude this discussion on the oligomeric forms of PKM2, it is important to look at the amino acids localized at the FBP-fixation pocket. While the monomer and dimer exhibit large structural pocket variations preventing it from any activity, the tetramer exhibits a bimodal distribution of the structures of the FBP-fixation pocket. This is observed for the free-tetramer where the reference Gly518 residue, which is chosen for the distance measurement, is localized in a well-known regulation-loop which has been experimentally shown to be highly flexible ^[37]^. Such flexibility is related to interactions of close amino acids including Trp515 and Arg516 that are implicated in the regulation of PKM2 being able to generate a less active tetramer known as the T-state and an alternative and more active form, the R-state. For the T-state, this loop is externalized from the FBP-fixation pocket and can form hydrogen bonds with the other chain nearby. This is also illustrated by our simulations where we showed higher distances between residues 433 and 518. On the opposite, for the R-state, the hydrogen bonds are lost since the loop that includes Gly518 is getting closer to the FBP fixation pocket exhibiting closer distances within the simulation. This is in agreement with previous structural data that compared different isoforms of PK where the rabbit structure of PKM1 do not possess a regulatory loop correctly orientated, thus blocking the fixation of FBP ^[18]^. The two tetramer states, observed in the simulations, are consistent with the experimentally observed equilibrium between the two states of the PKM2 tetramer, namely the R and T states ^[37]^. Taking into account every aspect of this comparison, our study provides good insight into the progressive structuration of PKM2 during the oligomerization process. In line with biological data ^[38]^, our results confirm its link with the overall enzyme activity as the PKM2 oligomerization progresses, having a particularly important function in the case of dimer/tetramer balance, could have implications in physiological cellular metabolism but also in the context of cancer. Indeed, It is important to remember that the predominance of the dimeric state of PKM2 in tumor cells is correlated with reduced enzymatic activity compared to tetramer but also the gain of nuclear functions by its possible transport in the nucleus, which promotes the accumulation of glycolysis metabolites, the use of alternate biosynthetic pathways but also the control of different genes for tumor development ^[39]^.

The second insight from our study is linked to the role of the natural FBP ligand in the structural regulation of PKM2 key pockets. First, the RMSD/RG profiles between the PKM2 apo and holo states exhibit a strong linear correlation in the holo-PKM2 condition. This is linked to the significant structural effect of FBP on the tetramer which is confirmed by the RMSF data. If they show very similar global profiles between both states, the holo state of PKM2 is nevertheless further stabilized in the C-terminal part of the protein that contains key residues for the fixation of FBP. It is important to note that even if FBP is docked into the PKM2 tetramer, the B-domain conserves its flexibility, especially in some small regions (Asn75, His78, Asp113, Thr114 and Arg120) implicated in the structuring of the active site. Such differences could explain the improved substrate accessibility for the active site through the allosteric effect of FBP. This is consistent with some previous theoretical results obtained on shorter molecular dynamics simulations on only one chain of two tetrameric crystal structures ^[40]^. The distance analysis of residues 53-129/178-296 localized at the active site validates this assumption. Overall, our results confirm that the fixation of FBP is important to maximize the structuration of the PKM2 active site thus allowing better enzymatic activity. Concerning the analysis of the FBP-fixation pocket itself, our results show the bimodal state of the tetramer PKM2 which reflects the existence of two different states, the T- and R-states. In the case of the holo-FBP form of PKM2, the residues that are constitutive of this pocket are less distant than in the apo state, an indication of structuring. Our results confirm previous works of Morgan and al. ^[10]^ concerning the effect of FBP on the R/T equilibrium of tetrameric PKM2 in human and parasites ^[41]^. This is further validated thanks to our modeling of a tetrameric state bound to the TEPP-46 pharmacological activator that is reminiscent of the one obtained with FBP. This is in line with biological data ^[19]^. Nevertheless, it is important to point out that the fixation region of TEPP-46 is localized in a different pocket than FBP inside the PKM2 structure and this could be a reason why the B-domain is more rigid in this case. However, TEPP-46 binding is still able to induce a satisfactory organization of the active site thanks to a stable binding after adding it to the structure. Interestingly, having a profile of the FBP-fixation pocket close to the PKM2+FBP condition even in the absence of FBP could explain the “FBP-like effect” of TEPP46 observed in the biological data ^[19]^. It is even more interesting since the stoichiometry of both compounds diverges by a factor of 2 with only two maximum pockets that can exist for TEPP-46 and 4 in the case of the natural allosteric ligand FBP, a plausible indication of a higher binding affinity of TEPP-46 compared to FBP.

Lastly, we also performed an initial characterization of the tetrameric state of PKM1, which is less well-described in the structural literature compared to PKM2. Our RMSD/RG analysis reveals the proximity of PKM1 to its PKM2 tetrameric counterpart, with or without FBP. PKM1 exhibits a profile that is intermediate between the holo and apo forms of PKM2. This is consistent with the fact that PKM1 is known to be a constitutively tetrameric protein that possesses enzymatic activity similar to that of FBP-PKM2 ^[42]^. Our RMSF data reveal the relative similarity of PKM1 to holo-PKM2, particularly in the B-domain and small key amino acids related to the active site. Nevertheless, some regions of PKM1 show higher RMSF flexibility compared to holo-PKM2. The first small region, more flexible in PKM1 compared to PKM2, is located at key residues 325-335 and includes the amino acid Thr328, which is implicated in the structuring of the PKM1/2 active site. Secondly, regions located at residues 470-490 and 515-525 both contain key amino acids involved in the PKM2 FBP-fixation pocket, where their flexibility is reduced. One hypothesis is that these unstable regions present in our PKM1 simulations could obstruct the formation of an effective FBP-pocket. Currently, no structural or experimental data document the effect of FBP on PKM1, except for some fixed data of the regulatory loop on an initial crystal structure comparing the FBP-fixation pockets of PKM1 and PKM2 obtained by Dombrauckas et al. ^[18]^. The last small region, located at residues 390-405, is specific to PKM1 since it is encoded by two different exons in PKM1 and PKM2. This result is surprising yet coherent since this region contains less flexible residues in the PKM2-specific exon (i.e., Tyr390, Gln393, and Leu394) that participate in the binding of known activators of PKM2 such as TEPP-46 and DASA-58 ^[43]^. One hypothesis could be that the relative instability of this key region is one of the reasons it is targeted in PKM2 but not in PKM1. To investigate the PKM1 active site, we studied the distance between amino acids present in both PKM isoforms. Our study aligns with biological data indicating that PKM1 is an active enzyme with a structured active site and where its B-domain maintains relative flexibility. This is consistent with results obtained for PKM2, as no exon-specific residues are implicated in the structuring of the active sites of both proteins ^[18]^. Moreover, there is the question of the existence (or absence) of the FBP-fixation pocket in PKM1, since no experimental data have shown any direct allosteric effects of FBP on PKM1. Our results lend further support to this hypothesis, as distances for the PKM1 tetrameric state are much greater than those in the PKM2 state bound to FBP. Unlike PKM2, which can exist in apo and holo-FBP states with two possible conformations, this is not the case for PKM1. The distribution of distances indicates a lack of stable structuration of a putative FBP-binding pocket in this region of PKM1. This data allows us to correlate the absence of an allosteric effect in PKM1 with the absence of an “allosteric pocket,” linked to exon-specific residues that are absent in the PKM1 isoform ^[6]^. The production of such precise models for both isoforms was crucial in helping us identify new cryptic pockets specific to PKM2 with potential for further drug design. Here, we discovered two new pockets that could impact the equilibrium between the T and R states of PKM2, one of which contains Cys423, close to its neighbor Cys424, implicated in the tetramerization of PKM2 ^[44]^. The other pocket, ‘T_M2-11’, contains Trp515, located in the regulatory loop previously shown in this study to be implicated in the FBP-fixation pocket. It is important to precise that the druggability scores of these pockets detected is a little low since usually one considers a pocket as “druggable” when its druggability score is higher than 0.5. Nevertheless, despite the extensive simulation effort, the identification of these pockets via molecular dynamics is based on a relatively small sample of structures that could be underestimated due to insufficient sampling. However, such results provide some hints for further experimental analysis.

## Conclusion

High-resolution *µ*s-scale molecular dynamics simulations were performed on both PKM1 and PKM2 enzymes using atomistic models. Whereas PKM1 is only expressed as an active tetramer, PKM2 models range from a monomer to a tetramer isoform. Such models allowed for an extensive study of the regulation of key domains such as the active site and the FBP allosteric pocket. First, at a global level, the PKM2 oligomerization process, which ranges from the monomer to the tetramer, shows a progressive PKM2 structuration while it remains capable to maintain some flexibility in its B-domain, which is intimately linked to the structure of the active site. In addition, despite the observation of small regions exhibiting higher flexibility, the PKM1 structural profile presents strong homology with the PKM2 tetramer bound to FBP. Moving to the FBP-fixation pocket, the latter is strongly influenced by the structural effect induced by the presence of the FBP ligand. This binding tends to select the R-state, which exhibits the most structured active site among the two states in equilibrium for this pocket when FBP is not present inside the tetramer. In parallel, the putative PKM1 FBP-fixation pocket tends to possess a larger volume compared to the FBP-pocket of PKM2, thus indicating an absence of FBP fixation while conserving a structured active site in agreement with biological data. The fixation to PKM2 of TEPP-46, a pharmacological activator that binds to a different pocket, was also analyzed and the structural differences/similarities compared to PKM2 bound to FBP discussed. Finally, a serie of PKM2-specific cryptic pockets were uncovered.

## Perspectives

Such a structural dynamics study on the different PKM1/2 isoforms adds further knowledge in the perspective of finding new molecules to modulate the enzymatic activity of the PKM enzyme. Indeed, such findings would be innovative in cancer therapy and also in the case of viral infections since PKM2 has been shown to participate in the viral replication of the Human Immunodeficiency Virus (HIV) ^[45]^ and of the pro-tumoral Epstein-Barr virus (EBV) ^[46]^. Only a small portion of ligands targeting PKM2, such as the synthetic TEPP-46 activator, has been tested in mouse models. Nevertheless, according to an initial study by Jiang et al. ^[47]^, no advanced clinical trials are presently ongoing even if some information concerning preclinical phases can still be found ^[19]^. Thus, further structural characterization and discovery of new druggable pockets targeting PKM2 remain of strong interest. In this context, different strategies to target PKM2 are possible using the computational models obtained in our study. These could go in three directions: i) targeting the FBP allosteric pocket; ii) targeting the TEPP interaction site; and iii) targeting other regions implicated in the PKM2 tetramerization. All these strategies converge towards a single goal: stabilizing the tetrameric state of PKM2, since the dimer has a positive effect on viral/tumor progression.

## Computational Details

### PDB Entries

The PKM1 system was constructed based on its apo tetrameric conformation obtained from the PDB entry 3SRF. We only considered this specific structure since it represents the only known active form of this enzyme. In contrast, PKM2 exhibits a broader range of oligomeric states. Conse-quently, we considered three initial systems for PKM2: the monomeric (PDB: 1ZJH), dimeric (PDB: 6B6U ^[48]^), and tetrameric (PDB: 3SRH) conformations, all in their apo states. To further investigate the impact of non–covalent interactions of the tetrameric PKM2 with various ligands, we also considered two holo forms of PKM2, associated with 1,6–di–O–phosphono–beta–D–fructofuranose (referred to as FBP, PDB: 3SRD) and TEPP-46 (PDB: 3U2Z ^[43]^), respectively.

### Preparation of the Systems

We selected six distinct systems, five corresponding to PKM2 and one to PKM1. For each of them, we removed all non–amino acid elements, retaining only ions, crystal waters, and for the two holo forms, their respective ligands. Subsequently, we prepared these six systems for molecular dynamics (MD) simulations using the AMOEBA (BIO18) polarizable force field ^[20,21]^. While parameters for amino acids as well as water and ions were readily available, standard parameters for accurately describing the dynamics of FBP and TEPP–46 ligands were not present in the literature. Thus, we generated them following the pipeline developed by Ren and al. ^[49]^ and employed the Poltype software ^[50]^. We ensured that the generated parameters provided stable MD simulations of each ligand by conducting short MD simulations in a water box. Once we assigned AMOEBA parameters to each molecule, periodic boundary conditions are applied via Smooth Particle Mesh Ewald ^[51]^ in conjunction with explicit solvation in a simulation box of length (100 Å x 100 Å x 100 Å for the monomer, 130 Å x 130 Å x 130 Å for the dimer, and 160 Å x 160 Å x 160 Å for the tetramers). We have thus six fully solvated initial structures of PKM2 monomer / dimer / tetramer-apo / tetramer+FBP / tetramer+TEPP-46 and PKM1 tetramer containing 95,267 / 212,664 / 394,651 / 394,679 / 395,732 and 394,828 atoms respectively. Subsequently, we modified the original pH of the structures obtained by crystallography thanks to the PDBFixer software by selecting physiological pH at 7.5 when choosing the protonation states, known to be optimal for PKM2 in the context of cancer cells ^[52]^.

### Conventional Molecular Dynamics

All MD simulations were performed using the Tinker-HP GPU software ^[27,53]^. Initially, each system underwent energy minimization using the L-BFGS algorithm with a Root Mean Square (RMS) threshold of 1 kcal/mol. Subsequently, the systems were gradually heated until equilibration was achieved. This was followed by 5 ns of NVT (canonical ensemble) Langevin dynamics simulations at 300K, using the BAOAB-RESPA1 integrator ^[54]^. Concerning the setup of the BAOAB-RESPA1 integrator, the bonded forces are always evaluated every 1 fs, but the short-range non-bonded ones are evaluated every 3.33fs, and the outer-timestep, i.e. the timestep of the long-range forces (that has to be a multiple of the previous one) is pushed up to 10fs using hydrogen mass repartitioning (HMR) ^[55]^. The Ewald cutoff parameter was set to 7 Å for Particle Mesh Ewald ^[51]^, and the van der Waals cutoff was set to a value of 9 Å. A convergence criterion of 10^*−*5^ was used for the self–consistent computations of the multipoles. These parameters remained consistent across all subsequent simulations. To further equilibrate the density, we conducted 5 ns of simulations in NPT (isothermal-isobaric ensemble), employing the same thermostat and temperature settings as before. Additionally, we used the Monte-Carlo barostat ^[56]^ to regulate the pressure at 1 Atmosphere. The last frame of each dynamic was taken as the starting point of the production stage, presented in the next section.

### Adaptive Sampling Simulations

The primary goal of this study was to capture the largest number of accessible conformations for each active conformation of PKM systems. To achieve this, we employed an enhanced sampling technique known as Adaptive Sampling (AS) ^[57]^ and based on the simulation protocol employed by Jaffrelot et al. in their previous work studying the Main Protease of SARS-CoV2 ^[28]^, which maximizes the exploration of rarely visited regions on the free energy surface associated to representative collective variables obtained through an unsupervised procedure. A notable advantage of this method lies in its unsupervised aspects, making it particularly versatile for studying diverse systems without a strong prior knowledge. AS studies were conducted for each of our six systems using the following pipeline.

An initial pool of structures was generated by running a 20 ns NPT simulation. Dimensionality reduction was then performed on all generated structures by computing the Principal Component Analysis (PCA) components using the sklearn library from Python ^[58]^. Subsequently, the structures were projected into a four-dimensional space using the first four PCA components, as they were found to capture over 90% of the structural diversity.

Following this initial phase, a series of 11 iterations was conducted. In each iteration, 10 structures were selected based on the same density law criterion used by Jaffrelot and al. ^[28]^, who demonstrated robustness in sampling for poorly visited regions. Subsequently, 9 ns of NPT simulations were performed on these 10 selected structures, resulting in a cumulative simulation time of 90 ns per AS iteration. The resulting structures were then added to the pool of conformations, and the process of structure selection was repeated until convergence in the conformational search was achieved.

By conducting 11 iterations, in addition to the initial 20 ns NPT simulation, the study ultimately provided a total simulation time of 1.01 *µ*s per system, amounting to a cumulative simulation time of 6.06 *µ*s for the entire study. The entire post–processing analysis were done using Python script with MDTraj ^[59]^, Numpy ^[60]^, Scikit–Learn ^[58]^ and Scipy ^[61]^ packages. Further graphical representations were done with Matplotlib ^[62]^ while the structural visualization were made with PyMOL (Version 3.0, Version 3.0 Schrödinger, LLC). Further detections of new pockets were performed with the DoGSite Scorer software ^[32]^ by using the default parameters on each selected PDB structures extracted from the molecular dynamics of the different PKM1 and PKM2 discussed oligomers.

## Acknowledgements

This work was made possible thanks to funding from both MESRI financing of the “Programme Doctorale Interdisciplinaire en Cancérologie (PDIC)” and the European Research Council (ERC) under the European Union’s Hori-zon 2020 research and innovation program (grant agreement Number 810367), project EMC2. Computations have been performed at GENCI (IDRIS, Orsay, France and TGCC, Bruyères le Chatel) on grant Number A0130712052. I would also like to give my thanks to M. Brandon Walker at the University of Texas at Austin for his help in the utilization of the Poltype software.

## Data availability

Simulation starting structures have been deposited on Zenodo repository: https://doi.org/10.5281/zenodo.14008130

## Code availability

TinkerHP is freely accessible to Academics via GitHub : https://github.com/TinkerTools/tinker-hp

Poltype is also freely accessible via Github : https://github.com/TinkerTools/poltype2/tree/master

## References

[1] B. Morten,H. M. R., K. P. E.G., van der Kogel Albert J., B. Johan, O. Jens, International Journal of Cancer 2008, 122, 2726.

[2] L. S. Y, V. H. M. G., Annu. Rev. Cell Dev. Biol. 2011, 27, 441.

[3] B. Emy, R. Claudio, Cells 2020, 9, 2073.

[4] I. Kiichi, T. Takehiko, The Journal of Biochemistry 1972, 6, 1043.

[5] N. Tamio, H. Inoue, T. Tanak, Journal of Biological Chemistry 1986, 261, 13807.

[6] W. J. Israelsen, M. G. V. Heiden, Seminars in cell & developmental biology 43 2015, pages 43–51.

[7] C. Barbara, D. Whitaker-Menezes, U. E. Martinez-Outschoorn, A. K. Witkiewicz, R. Birbe, A. Howell, R. G. Pestell, J. Smith, R. Daniel, F. Sotgia, M. P. Lisanti, Cancer biology & therapy 2011, 12, 1101.

[8] C. V. Clover, D. Chatterjee, Z. Wang, L. C. Cantley, M. G. V. Heiden, A. R. Krainer, Proceedings of the National Academy of Sciences of the United States of America 2010, 5, 1894.

[9] M. Takenaka, N. Tamio, S. Shigeki, H. Haruhiko, Y. Kazuya, M. Tamiko, I. Enyu, T. Takehiko, European Journal of Biochemistry 1991, 1, 101.

[10] H. P. Morgan, F. J. O’Reilly, M. A. Wear, J. R. O’Neill, L. A. Fothergill-Gilmore, T. Hupp, M. D. Walkinshaw, Proceedings of the National Academy of Sciences 110 2013, 15, 5881.

[11] Y. Weiwei, Z. Lu., Cell cycle 2013, 12,19, 3154.

[12] Y. Ikeda, T. Tanaka, T. Noguchi, Journal of Biological Chemistry 272 1997, 33, 20495.

[13] M. Yuan, M. I. W., C. Yiyuan, B. E. A., W. M. A., M. P. A.M., F.-G. L. A., H. Ted, W. M. D., Biochemical Journal 475 2018, 10, 1821.

[14] K. E. Keller, I. S. Tan, Y.-S. Lee, Science (New York, N.Y.) 338 2012, 6110, 1069.

[15] M. S. Jurica, A. Mesecar, P. J. Heath, B. L. Stoddard, W. Shi, T. Nowak, Structure 6 1998, 2, 195.

[16] H. Kato, T. Fukuda, C. Parkison, P. McPhie, S.-Y. Cheng, Proceedings of the National Academy of Sciences of the United States of America 86 1989, 20, 7861.

[17] G. Gdynia, S. W. Sauer, J. Kopitz, D. Fuchs, K. Duglova, T. Ruppert, M. Miller, J. Pahl, A. Cerwenka, M. Enders, H. Mairbäurl, M. M. Kamiński, R. Penzel, C. Zhang, J. C. Fuller, R. C. Wade, A. Benner, J. Chang-Claude, H. Brenner, M. Hoffmeister, H. Zentgraf, P. Schirmacher, W. Roth, Nature Communications 2016, 7, 10761.

[18] J. D. Dombrauckas, B. D. Santarsiero, A. D. Mesecar, Biochemistry 44 2005, 27, 9417.

[19] D. Anastasiou, Y. Yu, W. J. Israelsen, J.-K. Jiang, M. B. Boxer, B. S. Hong, W. Tempel, S. Dimov, M. Shen, A. Jha, H. Yang, K. R. Mattaini, C. M. Metallo, B. P. Fiske, K. D. Courtney, S. Malstrom, T. M. K. and Charles Kung, A. P. Skoumbourdis, H. Veith, N. Southall, M. J. Walsh, K. R. Brimacombe, W. L. and Sophia Y Lunt, Z. R. Johnson, K. E. Yen, K. Kunii, S. M. Davidson, H. R. Christofk, C. P. Austin, J. Inglese, M. H. Harris, J. M. Asara, G. Stephanopoulos, F. G. Salituro, S. Jin, L. Dang, D. S. Auld, H.-W. Park, L. C. Cantley, C. J. Thomas, M. G. V. Heiden, Nature chemical biology 8 2012, 10, 839.

[20] P. Ren, J. W. Ponder, The Journal of Physical Chemistry B 107 2003, 24, 5933.

[21] Y. Shi, Z. Xia, J. Zhang, R. Best, C. Wu, J. W. Ponder, P. Ren, Journal of chemical theory and computation 9, 2013, 9, 4046.

[22] N. Gresh, G. A. Cisneros, T. A. Darden, J.-P. Piquemal, Journal of Chemical Theory and Computation 2007, 3, 1960, pMID: 18978934.

[23] Y. Shi, P. Ren, M. Schnieders, J.-P. Piquemal, Polarizable Force Fields for Biomolecular Modeling, chapter 2, pages 51–86, John Wiley & Sons, Ltd 2015.

[24] J. Melcr, J.-P. Piquemal, Frontiers in Molecular Biosciences 2019, 6, 143.

[25] Z. Jing, C. Liu, S. Y. Cheng, R. Qi, B. D. Walker, J.-P. Piquemal, P. Ren, Annual Review of Biophysics 2019, 48, 371, pMID: 30916997.

[26] O. Adjoua, L. Lagardère, L.-H. Jolly, A. Durocher, T. Very, I. Dupays, Z. Wang, T. J. Inizan, F. Célerse, P. Ren, J. W. Ponder, J.-P. Piquemal, Journal of Chemical Theory and Computation 17 2021, 4, 2034.

[27] L. Lagardère, J. Luc-Henri, L. Filippo, A. Félix, S. Benjamin, J. Z. F., H. Matthew, T. Hedieh, C. G. Andrés, S. M. J., G. Nohad, M. Yvon, R. P. Y., P. J. W., P. Jean-Philip, Chemical Science 9 2018, 4, 956.

[28] T. J. Inizan, C. Frédéric, A. Olivier, E. A. Dina, J. Luc-Henri, L. Chengwen, R. Pengyu, M. Matthieu, L. Nathalie, L. Louis, M. Pierre, P. Jean-Philip, Chemical Science 12 2021, 13, 4889.

[29] E. A. Dina, L. Lagardère, T. J. Inizan, F. Célerse, C. Liu, O. Adjoua, L.-H. Jolly, N. Gresh, Z. Hobaika, P. Ren, R. G. Maroun, J.-P. Piquemal, J. Phys. Chem. Lett. 2021, 12, 6218.

[30] L. El Khoury, Z. Jing, A. Cuzzolin, A. Deplano, D. Loco, B. Sattarov, F. Hédin, S. Wendeborn, C. Ho, D. El Ahdab, T. Jaffrelot Inizan, M. Sturlese, A. Sosic, M. Volpiana, A. Lugato, M. Barone, B. Gatto, M. L. Macchia, M. Bellanda, R. Battistutta, C. Salata, I. Kondratov, R. Iminov, A. Khairulin, Y. Mykhalonok, A. Pochepko, V. Chashka-Ratushnyi, I. Kos, S. Moro, M. Montes, P. Ren, J. W. Ponder, L. Lagardère, J.-P. Piquemal, D. Sabbadin, Chem. Sci. 2022, 13, 3674.

[31] M. Blazhynska, L. Lagardère, C. Liu, O. Adjoua, P. Ren, J.-P. Piquemal, Chem. Sci. 2024, 15, 14177.

[32] V. Andrea, K. Daniel, R. Friedrich, R. Matthias, Bioinformatics 2012, 28, 2074.

[33] P. Kalaiarasan, N. Subbarao, R. Bamezai, Journal of molecular modeling 20 2014, page 2447.

[34] Z. Zhang, X. Deng, Y. Liu, Y. Liu, L. Sun, F. Chen, Cell & Bioscience 9 2019, 1, 52.

[35] X. Jiansheng, C. Dai, X. Hu, The Journal of Biological Chemistry 291 2016, 17, 8987.

[36] G. Jianshuand, X. Qingqing, L. Kaihui, G. Weizhi, L. Wenjie, W. Jiyan, Z. Mengyi, L. Qiu-ying, C. Dongpo, S. Changliang, Z. Chunze, L. Xinqi, L. Jing, Frontiers in Oncology 2019, 9, 993.

[37] W. Ping, C. Sun, T. Zhu, Y. Xu, Protein & cell 6 2015.

[38] G. Xueliang, W. Haizhen, Y. J. J, L. Xiaowei, L. Zhi-Ren, Molecular Cell 45 2012, 5, 598.

[39] K. Zahra, D. Tulika, Ashish, M. S. Pratap, P. Uma, Frontiers in Oncology 2020, 10, 2234.

[40] Y. Jingxu, H. Liu, X. Liu, C. Gu, R. Luo, H.-F. Chen, Journal of chemical information and modeling 56 2016, 6, 1184.

[41] M. H. P., I. W. McNae, M. W. Nowicki, V. Hannaert, P. A. Michels, L. A. Fothergill-Gilmore, M. D. Walkinshaw, Journal of Biological Chemistry 285 2010, 17, 12892.

[42] C. Heather, M. G. V. Heiden, N. Wu, J. M. Asara, L. C. Cantley, Nature 452 2008, pages 181–86.

[43] A. Dimitrios, Y. Yu, W. J. Israelsen, J.-K. Jiang, M. B. Boxer, B. S. Hong, W. Tempel, S. Dimov, M. Shen, A. Jha, H. Yang, K. R. Mattaini, C. M. Metallo, B. P. Fiske, K. D. Courtney, S. Malstrom, T. M. Khan, C. Kung, A. P. Skoumbourdis, H. Veith, N. Southall, M. J. Walsh, K. R. Brimacombe, W. Leister, S. Y. Lunt, Z. R. Johnson, K. E. Yen, K. Kunii, S. M. Davidson, H. R. Christofk, C. P. Austin, J. Inglese, M. H. Harris, J. M. Asara, G. Stephanopoulos, F. G. Salituro, S. Jin, L. Dang, D. S. Auld, H.-W. Park, L. C. Cantley, C. J. Thomas, M. G. V. Heiden, Nature chemical biology 8 2012, 10, 839.

[44] M. So, K. Hashimoto, D. Kihara, C. Tsuzuki, N. Kataoka, K. Suzuki, Biochemical and Biophysical Research Communications 2020, 526, 973.

[45] Z. Hong-Sheng, Z.-G. Zhang, Z. Zhou, G.-Y. Du, H. Li, X.-Y. Yu, Y.-H. Huang, Biochemistry and Biophysics 2017, 625-626, 17–23.

[46] L. A. Kwok-Fung, D. C. W, Y. L. S, K. Chuen-Wai, H. Pok-Man, L. Kwok-Wai, The Journal of Pathology 2015, 237, 238.

[47] J. Jian-kang, M. B. Boxer, M. G. Vander Heiden, M. Shen, A. P. Skoumbourdis, N. Southall, H. Veith, W. Leister, C. P. Austin, H. W. Park, J. Inglese, L. C. Cantley, D. S. Auld, C. J. Thomas, Bioorganic medicinal chemistry letters 2010, 20, 3387.

[48] D. Srivastava, M. Razzaghi, M. T. Henzl, M. Dey, Biochemistry 2017, 2017, 6517.

[49] J. C. Wu, G. Chattree, P. Ren, Theor Chem Acc 2012, 131, 1138.

[50] B. Walker, L. Chengwen, W. Elizabeth, R. Pengyu, Journal of Computational Chemistry 2022, 43, 1530.

[51] L. Lagardère, F. Lipparini, Polack, B. Stamm, Cancès, M. Schnieders, P. Ren, Y. Maday, J.-P. Piquemal, Journal of Chemical Theory and Computation 2015, 11, 2589, pMID: 26575557.

[52] S. Nandi, R. Mortezaali, S. Dhiraj, D. Mishtu, Journal of Biological Chemistry 295 2020, 51, 17425.

[53] R. Joshua, Z. Wang, C. Lu, M. L. Laury, L. Lagardère, M. J. Schnieders, J.-P. Piquemal, P. Ren, J. W. Ponder, J. Chem. Theory Comput. 2018, 14, 5273.

[54] L. Lagardère, F. Aviat, J.-P. Piquemal, J. Phys. Chem. Lett. 2019, 10, 2593.

[55] H. C. W, S. L. Grand, R. C. Walker, A. E. Roitberg, J. Chem. Theory Comput. 2015, 11, 1864.

[56] J. Åqvist, P. Wennerström, M. Nervall, S. Bjelic, B. O. Brandsdal, Chemical Physics Letters 2004, 384, 288.

[57] G. R. Bowman, D. L. Ensign, V. S. Pande, J. Chem. Theory Comput. 2010, 6, 787.

[58] F. Pedregosa, G. Varoquaux, A. Gramfort, V. Michel, B. Thirion, O. Grisel, M. Blondel, A. Müller, J. Nothman, G. Louppe, P. Prettenhofer, R. Weiss, V. Dubourg, J. Vanderplas, A. Passos, D. Cournapeau, M. Brucher, M. Perrot, Édouard Duchesnay, arXiv 2018.

[59] R. T. McGibbon, B. K. A, H. M. P, K. Christoph, S. J. M, H. C. X, S. C. R, W. Lee-Ping, L. T. J, P. V. S, Biophysical Journal 2015, 8, 1528.

[60] C. R. Harris, K. J. Millman, S. J. van der Walt, R. Gommers, P. Virtanen, D. Cournapeau, E. Wieser, J. Taylor, S. Berg, N. J. Smith, R. Kern, M. Picus, S. Hoyer, M. H. van Kerkwijk, M. Brett, A. Haldane, J. F. del Río, M. Wiebe, P. Peterson, P. Gérard-Marchant, K. Sheppard, T. Reddy, W. Weckesser, H. Abbasi, C. Gohlke, T. E. Oliphant, Nature 2020, 7825, 357.

[61] P. Virtanen, R. Gommers, T. E. Oliphant, M. Haberland, T. Reddy, D. Cournapeau, E. Burovski, P. Peterson, W. Weckesser, J. Bright, S. J. van der Walt, M. Brett, J. Wilson, K. J. Millman, N. Mayorov, A. R. J. Nelson, E. Jones, R. Kern, E. Larson, C. J. Carey, İlhan Polat, Y. Feng, E. W. Moore, J. VanderPlas, D. Laxalde, J. Perktold, R. Cimrman, I. Henriksen, E. A. Quintero, C. R. Harris, A. M. Archibald, A. H. Ribeiro, F. Pedregosa, P. van Mulbregt, S. Contributors, Nature Methods 2020, 3, 261.

[62] J. D. Hunter, Computing in Science & Engineering 2007, 3, 90.

