## Supplementary Information for "High-Resolution Molecular-Dynamics Simulations of the Pyruvate Kinase Muscle Isoform 1 and 2 (PKM1/2)"

### Supplementary Figures

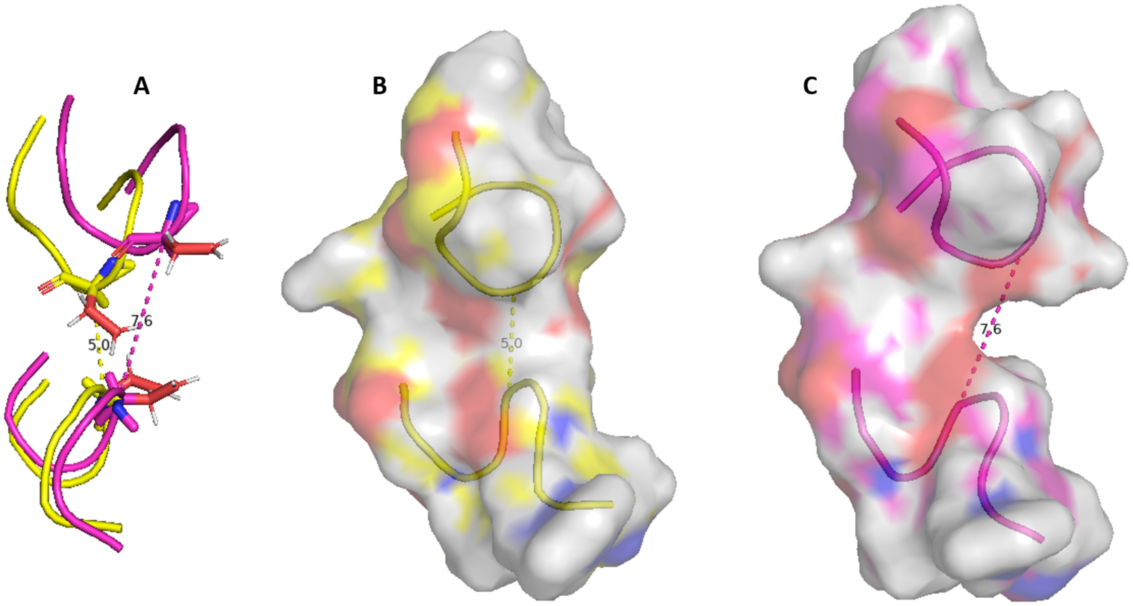


Figure S1: Comparative study of different states of the active side internal part of dimeric PKM2. (A) Graphical representation of the distance between residues 178 and 296 for two different frames of the bimodal distribution, **(B)** Surface model showing the low-distance state ($\approx 5$ Å) **(C)** or the largest one ($\approx 7.5$ Å). Three amino acids before and after residues 178 and 296 are represented.

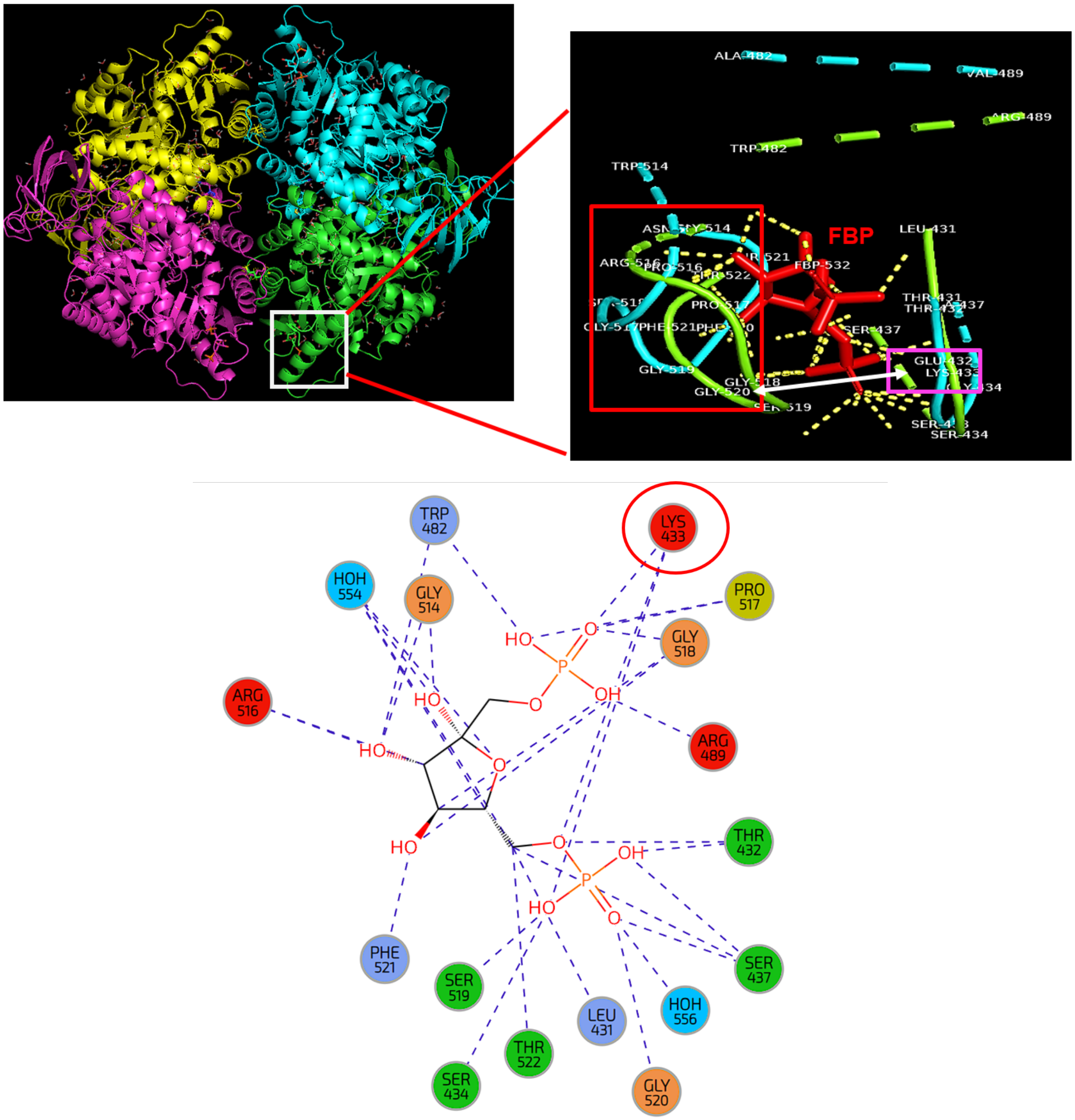


Figure S2: Structural representation of the FBP-binding site of PKM isoforms. Cartoon model of the FBP-binding site and comparison between PKM1 (in blue) and PKM2 (in green) in presence of FBP (in red). Important sites are colored to highlight their importance in the binding of FBP: the red square represents a regulatory loop and the violet square the residue 433 (Lys for PKM2 and Glu for PKM1). Further below are represented the binding environment of FBP with the corresponding PKM2 residues from the PDB crystal structure.

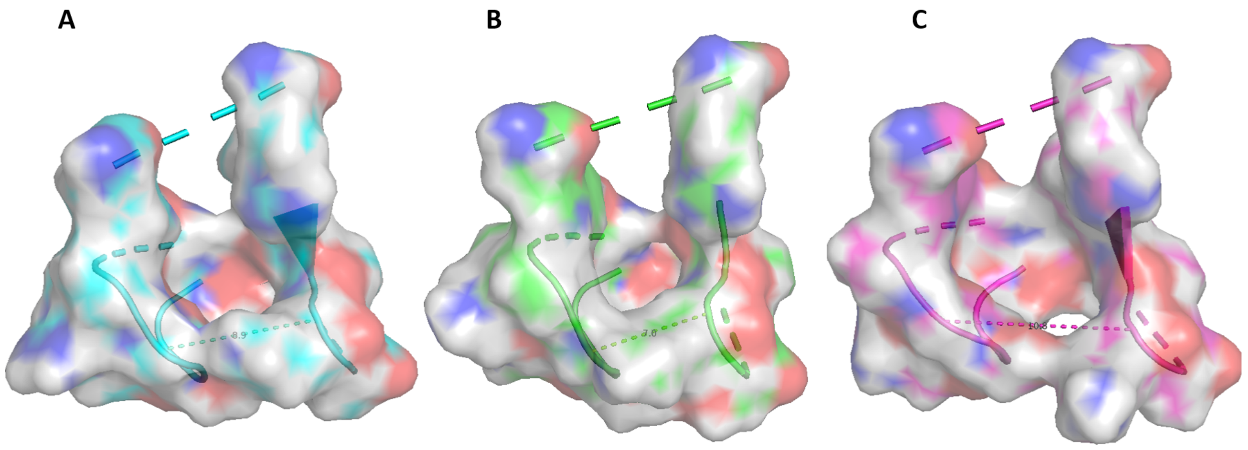


Figure S3: Comparative study of the different states of the FBP-fixation pocket for PKM2 bound to FBP. Surface model of three frames aligned during simulation of PKM2 bound to FBP with the lowest (≈ 7 Å) **(A)** medium (≈ 9 Å) **(B)** and highest (≈ 11 Å) **(C)** distances between residues 433 and 518.

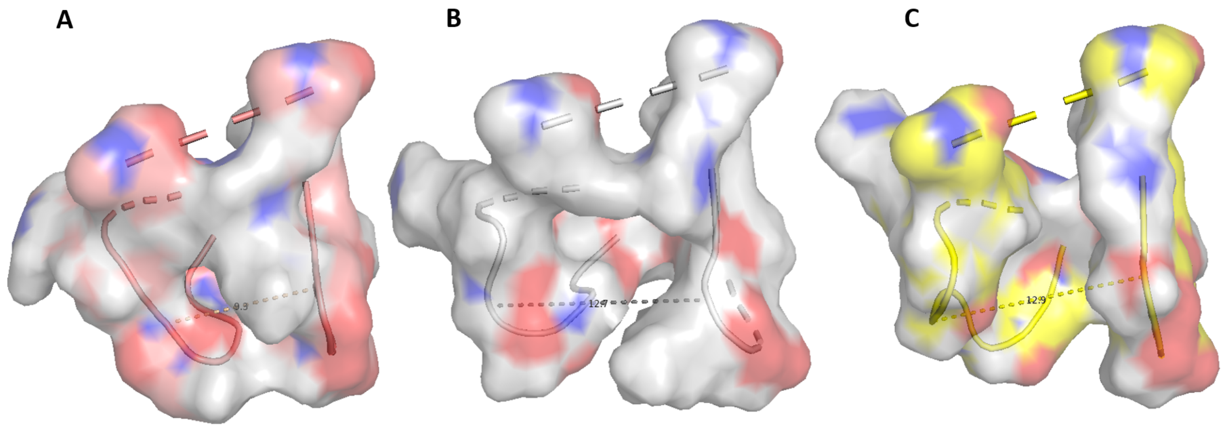


Figure S4: Comparative study of the different states of the FBP-fixation pocket for free-PKM2 and PKM1. Surface model of two frames aligned during simulation of PKM2 not bound to FBP with **(A)** the lowest (≈ 9 Å) and **(B)** the highest (≈ 12, 5 Å) distances from the bimodal state between residues 433 and 518. The same model is represented for the tetrameric state of PKM1 (≈ 13 Å) **(C)**.

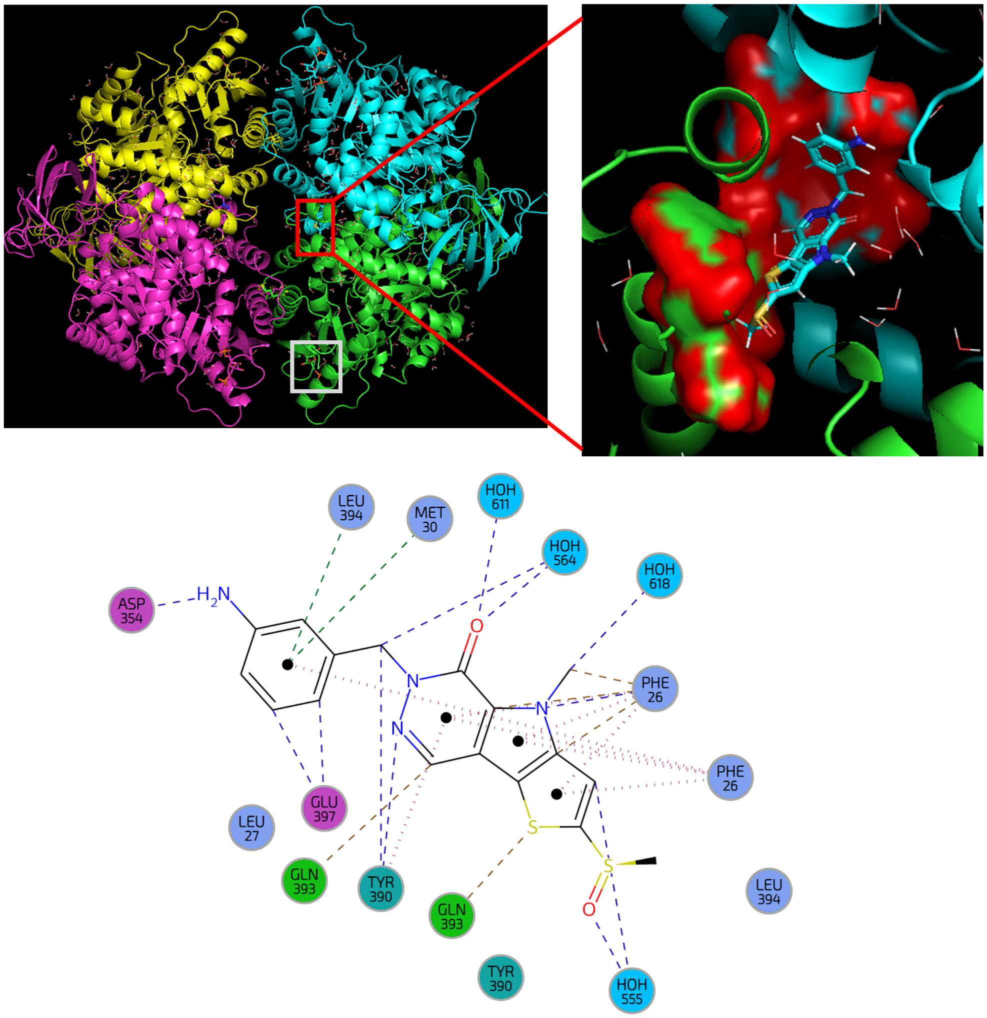


Figure S5: Structural representation of the TEPP-46 fixation site on PKM2. Cartoon model of PKM2. The red frame indicates the TEPP-46 binding regions. The white frame indicates the FBP binding regions. Several residues that interact with TEPP-46 are shown in surface format and are colored in red. Further below are represented the binding environment of TEPP-46 with the corresponding PKM2 residues in the PDB crystal structure.

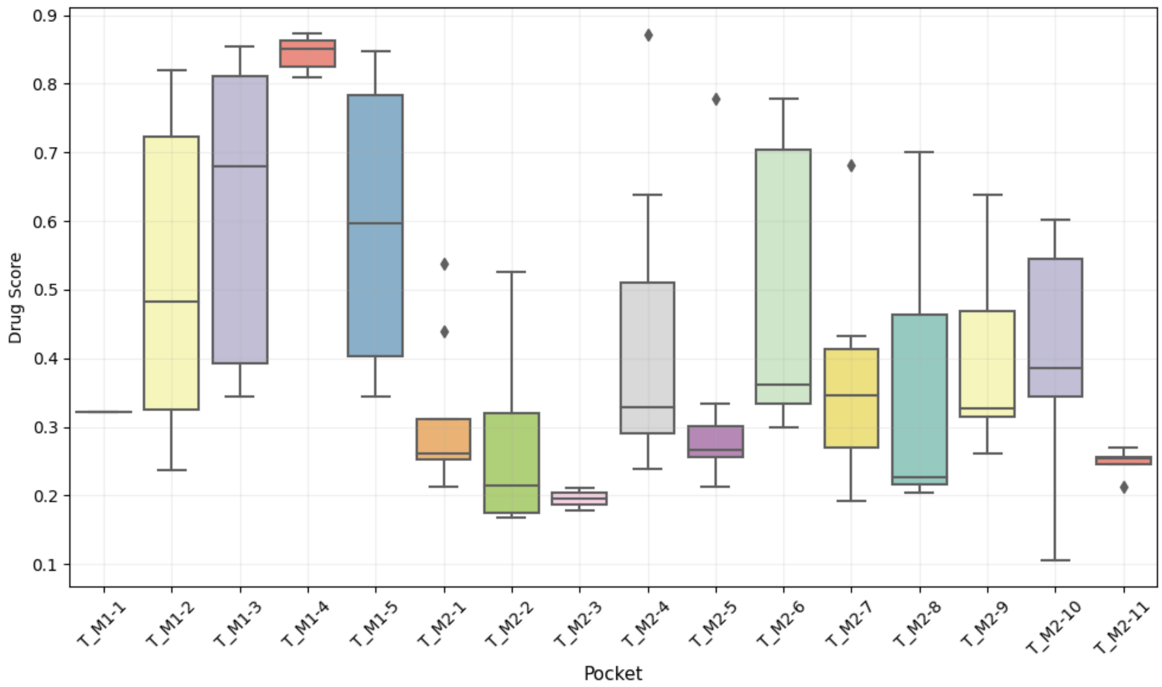


Figure S6: Boxplot of DrugScore distribution for each new pockets obtained by structural sampling. The new pockets were found from the selection of some structures from the molecular dynamics of PKM1 (M1-X) or PKM2 (M2-X) that weren't found on the initial structures 3SRF (tetrameric cristal of PKM1) and 3SRD (tetrameric state of PKM2 bound to FBP).

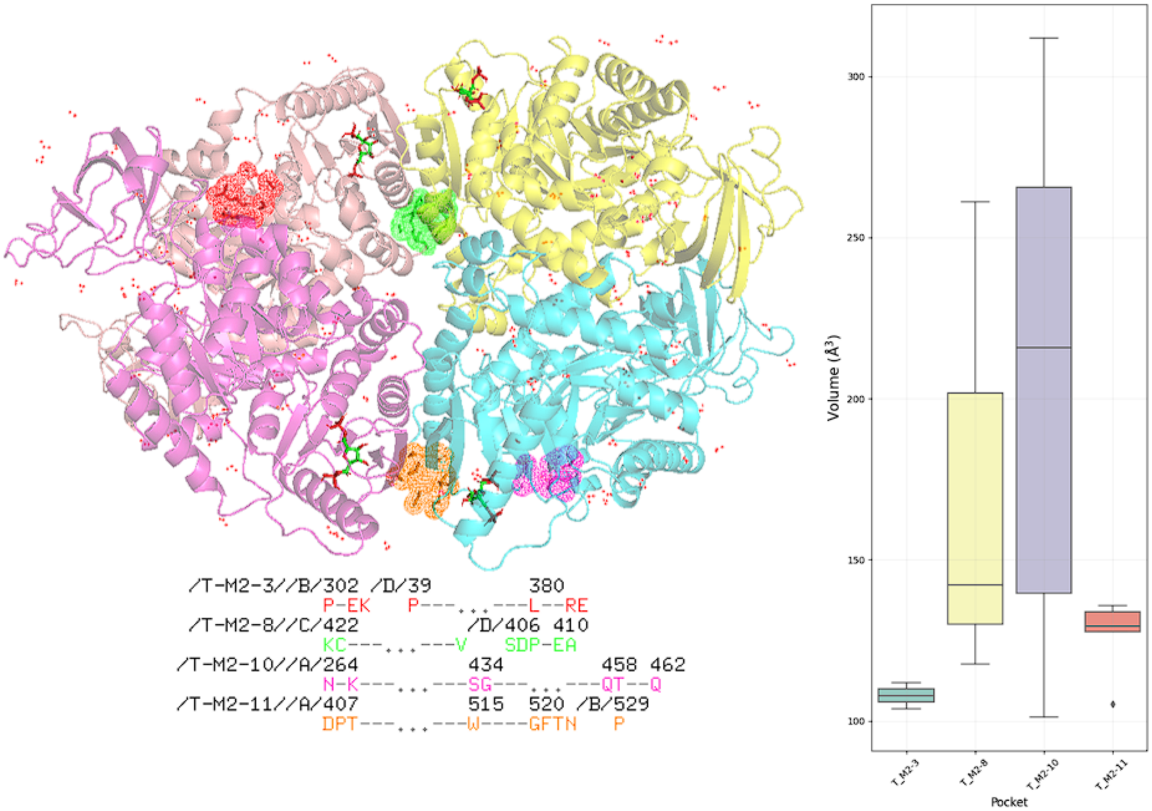


Figure S7: Structures and volumes of potential new pockets specific for PKM2 determined by a small sampling of the Tinker-HP simulations. The residues implicated in pockets structures are shown on the left panel in red, green, orange and pink with structural visualization on the initial crystal structure. The volumes of each cavity is shown on the right panel with a box-plot representation.

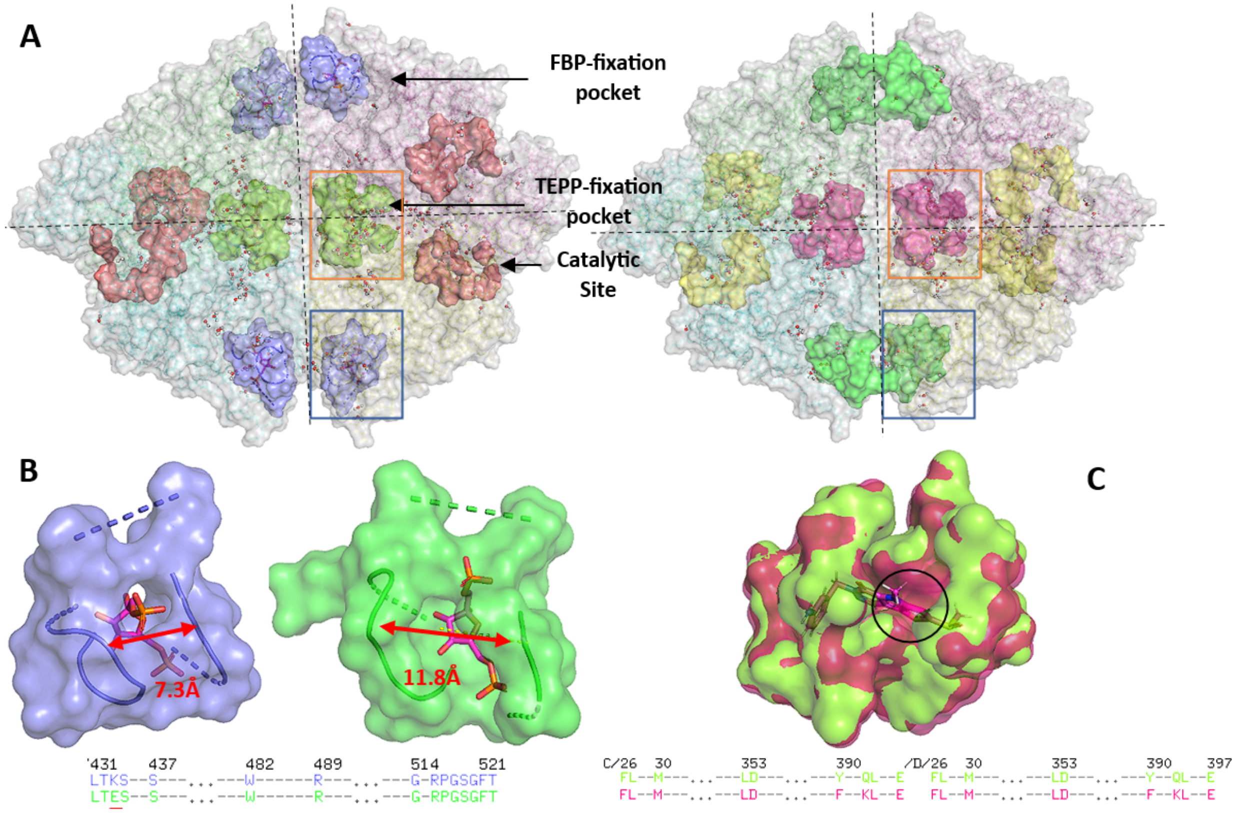


Figure S8: Global view of each PKM enzymes with their major site. (A) Surface model of PKM2 (left side) and PKM1 (right side) associated with key residues corresponding to region structuring the FBP site (light blue (PKM2) and green (PKM1)), TEPP site (light green (PKM2) and red (PKM1)) and the catalytic site (light red (PKM2) and yellow (PKM1)). Water molecules localized at 9 Å or less are represented in CPK model. **(B)** Detailed surface model of the residues implicated in FBP fixation for PKM2 (left) and same view for PKM1. The red arrow corresponds to the distance between residues 433 and 518. **(C)** Surface model of the residues interacting with TEPP-46 for the comparison between PKM2 (light green) and PKM1 (red). One major difference is localized at residue 26 for PKM1 with a more closed pocket compared to PKM2 (black circle).
